# A biobank-scale method for learning modulators of gene-environment interaction underlying human complex traits from multiple environmental exposures

**DOI:** 10.64898/2026.03.13.711725

**Authors:** Zhengtong Liu, Arush Ramteke, Aakarsh Anand, Aditya Gorla, Moonseong Jeong, Sriram Sankararaman

## Abstract

It is increasingly recognized that genetic effects on complex traits and diseases are shaped by environmental context. Biobanks that measure diverse environmental exposures alongside genotypes and phenotypes at scale enable systematic study of gene-environment (G×E) interactions. Existing approaches, however, are limited in their ability to accurately model polygenic G×E involving many exposures across genome-wide genetic variants. It is unclear which exposure combinations are relevant for a given trait while distinguishing true interactions from environment-dependent heteroskedastic noise. To address these challenges, we develop *Efficient multi-eNvironmental Gene-environment Interaction iNference Estimator* (ENGINE), a supervised variance-component framework that learns an embedding that combines multiple environmental exposures while jointly estimating additive, G×E, and heteroskedastic noise components. To enable biobank-scale inference, ENGINE makes a single pass over the genotype matrix to cache genotype-dependent summaries, then assembles normal-equation components and gradients at each iteration. In simulations, ENGINE controls type I error rates, achieves high power, and accurately recovers the environmental embedding while remaining efficient at biobank-scale. Applied to five complex traits paired with lifestyle exposures in *N* = 291,273 unrelated white British individuals and *M* = 454,207 common SNPs (MAF> 0.01) from the UK Biobank, ENGINE recovered G×E variance that was on average 1.4-fold larger than that captured by a single exposure and 5.5-fold larger than that captured by the first principal component of the exposures.

## 1 Introduction

Gene–environment (G×E) interactions—wherein the magnitude and sometimes the direction of a variant’s effect depends on environmental context—are increasingly recognized as important contributors to the genetic architecture of complex traits [1–5]. Growing empirical evidence for context-dependent allelic effects across environments [6, 7] suggests that explicitly modeling G×E can increase explained variance, help account for components of missing heritability, and improve risk stratification in predictive models [8–12].

In practice, the operative “environment” is rarely a single exposure [13–17]. Large biobanks now measure a wide array of exposures that collectively constitute the exposome [18–22]. Within this exposome, lifestyle and behavioral factors, medication use, and sociodemographic context covary and can jointly modulate genetic effects via G×E. For instance, low-density lipoprotein (LDL) levels may reflect interactions between lipid-related loci and age, while statin use, diet, and physical activity further modify those genetic influences [8, 23–25]. Modeling such multi-environment modulation requires methods that can accommodate many environmental variables, retain interpretability at the level of individual environments, and scale to biobank-sized cohorts.

Polygenic models quantify the genome-wide contribution of G×E by aggregating variant-level interaction effects, thereby offering higher power than single-variant approaches when individual effects are weak. In particular, variance component or mixed models are well-suited to model aggregate genetic signal, arising from additive genetic effects and from interactions such as those due to G × E, within a single framework [7, 26–29]. Early variance component approaches for modeling polygenic G × E (e.g., GCTA-G × E [30], MV-GREML [31], MRNM [26]) enabled inference for interactions with individual environmental exposures. LEMMA [32] models polygenic G × E by learning an environment score—a data-driven combination of exposures–that allows multiple environments to jointly modulate genetic effects. However, because LEMMA does not explicitly model environment-dependent residual variance, estimates of G × E variance can be biased upward under noise heterogeneity, leading to inflated false positive rates [8]. MonsterLM [9] scales to biobank-sized datasets and yields approximately unbiased estimates in many settings; however, its reliance on pruning and partitioning the genome into relatively weakly correlated blocks (and its focus on common variation) can limit power and scope for discovering G × E effects. GxEMM [7] and GENIE [8] incorporate context-specific heteroskedasticity within a variance-components framework; GENIE is designed to scale to biobank-sized cohorts, whereas GxEMM becomes computationally infeasible for large samples. Despite these advances, approaches to learn which combinations of environmental exposures contribute to polygenic G × E on a biobank-scale remain limited.

Here we present ENGINE, *Efficient multi-eNvironmental Gene-environment Interaction iNference Estimator*, a method that jointly learns the combination of environments that interact with polygenic effects for a given trait, estimates the contribution of each environment to the interaction signal while producing calibrated variance-component estimates for additive effects, G × E, and noise by explicitly modeling environment-dependent heteroskedasticity (Figure 1). ENGINE scales to biobank-sized cohorts by caching genotype-dependent intermediate quantities so that subsequent model updates avoid repeated full passes over the genotype matrix. In simulations, ENGINE is well calibrated and accurately estimates the variance components and the environmental embedding. Applied to five complex traits with lifestyle exposures in UK Biobank, ENGINE detects substantially larger G × E signal than analyses that consider environments one at a time or replace them with unsupervised embeddings. Together, ENGINE provides a principled and tractable path to disentangling how multiple environments jointly modulate genetic effects on complex traits.

**Figure 1.**
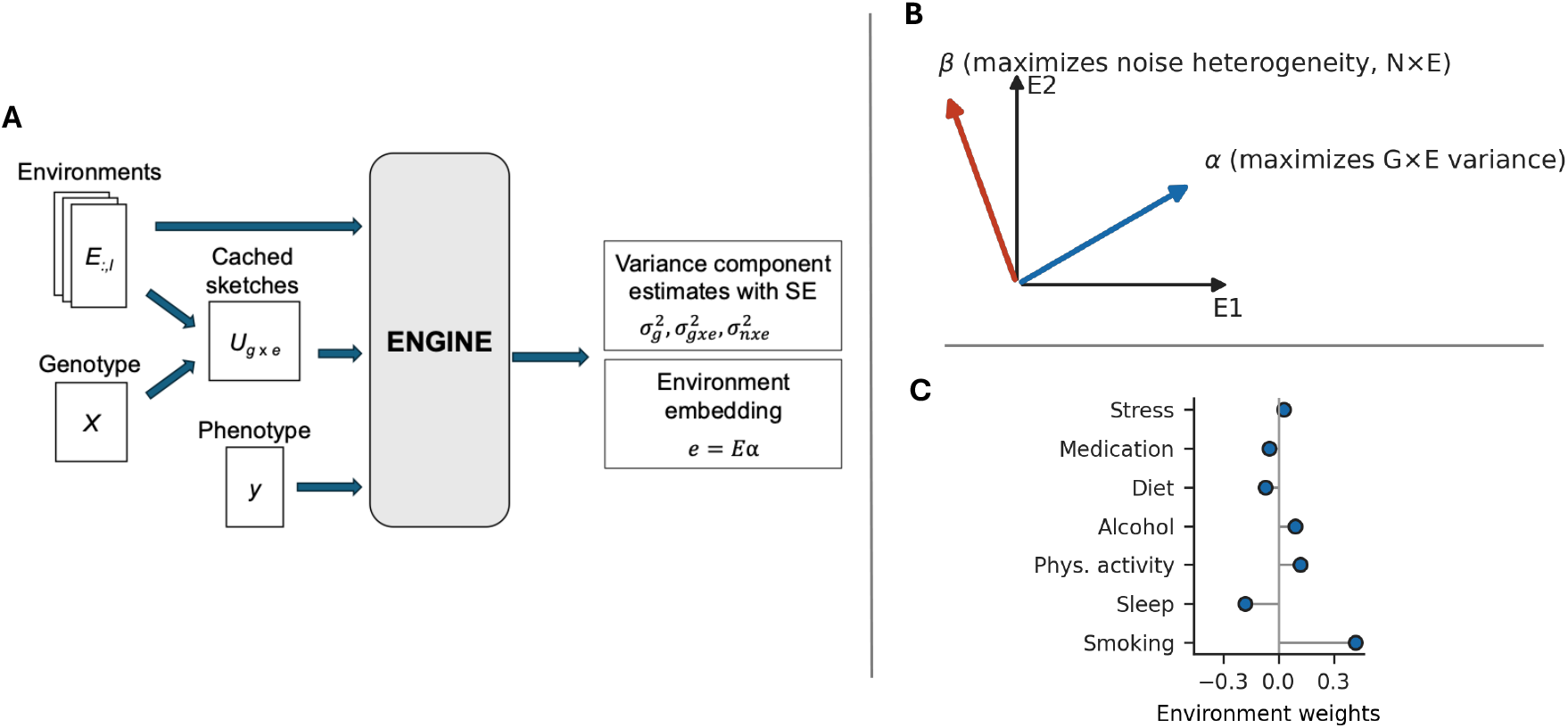
Overview of ENGINE. **A. Pipeline.** Inputs are a genotype matrix, multiple environments, and a phenotype. ENGINE learns an interpretable environmental embedding ***e*** and reports point estimates and standard errors for the additive 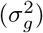, interaction 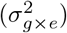, and noise-by-environment 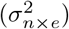 variance components. For efficiency, genotype-dependent summaries (*U*_*g*×*e*_) are cached in a single streaming pass and reused across iterations. **B. Separating heteroskedastic noise from G** × **E**. With two environments, the combination that maximizes N × E variance can differ from the one that maximizes G × E variance; ENGINE therefore learns weights that maximize G × E while controlling N × E. **C. Interpretable environment weights**. The entries of environmental weights (***α***) quantify the contribution of each environmental exposure to the embedding and to the captured G×E signal.

## 2 Methods

### 2.1 Estimation of variance components in G×E linear mixed model

Let ***X*** ∈ ℝ^*N*×*M*^ denote standardized genotypes of *M* variants on *N* individuals, ***y*** ∈ ℝ^*N*^ a standardized phenotype, and ***E*** ∈ ℝ^*N*×*L*^ a matrix of *L* measured environments (columns are centered and scaled). We compress the *L* environments into a one–dimensional vector ***e*** = ***Eα***, with interpretable, unit–norm weights ***α*** ∈ ℝ^*L*^ that quantify each environment’s contribution to the interaction signal. Intuitively, ***e*** collects heterogeneous environmental effects into a single direction along which genetic effects may be modulated.

We model the phenotype ***y*** with a linear mixed model comprising four components: an additive genetic effect, a G ×E interaction that varies with the environment embedding ***e***, a noise-by-environment (N × E) term that captures heteroskedasticity aligned with ***e***, and a residual term [29, 28, 33, 7]:

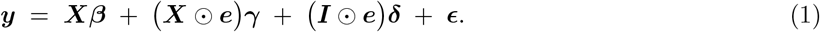

Here, ⊙ denotes the Hadamard (elementwise) product with row-wise broadcasting across SNP columns, so that ***X*** ⊙ ***e*** = diag (***e***) ***X*** and ***I*** ⊙ ***e*** = diag (***e***); equivalently, we scale each individual genotype and residual noise by their environmental score. The N × E component (context-specific noise) captures heteroskedastic noise across individuals exposed to different environments. Distinguishing this component from the G × E component is crucial for controlling false positives, and such environment-dependent heteroskedasticity is widely observed for complex traits [8].

We assume independent, mean-zero random effects with variances that define the components of interest:

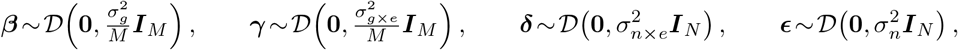

where 𝒟 can be any zero-mean distribution with the stated second moments (this moment assumption makes the method-of-moments robust to non-Gaussian effect distributions). The M-vectors ***β*** and ***γ*** represent the main additive effects and the G× E interaction effects along ***e*** respectively, while ***δ*** captures environment-modulated noise (Table S1). Under 𝔼 [***y***] = **0**, the population covariance decomposes additively into kernels:

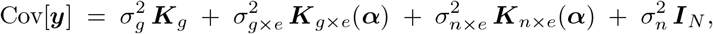

where

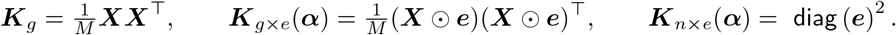

Here ***K***_*g*_ is the genetic relatedness matrix (GRM), ***K***_*g*×*e*_ is the GRM after scaling each individual’s genotypes by *e*_*i*_, thereby up-weighting pairs of individuals who are both environmentally exposed, and ***K***_*n*×*e*_ is diagonal with entries 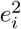, modeling environment-dependent variance inflation.

Given fixed ***α***, we first estimate the variance components 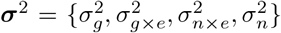 using the method-of-moments (MoM), matching the model covariance to the empirical covariance ***yy***^⊤^ via a Frobenius-norm objective:

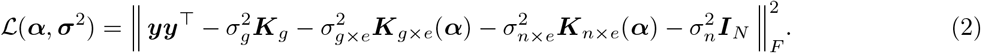

Because Eq. (2) is quadratic in ***σ***^**2**^, the minimizer 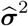 solves the linear system 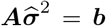, where ***A*** ∈ ℝ^4×4^, ***b*** ∈ ℝ^4^ are defined:

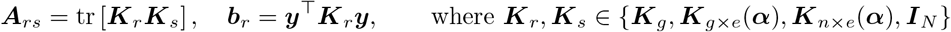

We attach asymptotic standard errors to 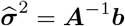 via the delta method:

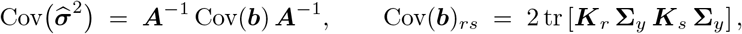

with **Σ**_*y*_ = Cov[***y***]. In practice, we plug in 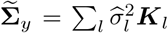 and evaluate the trace contractions efficiently using probe sketches (see Section 2.3). Because ***A*** is small and reused, SE computation adds negligible 106 overhead once the necessary contractions are cached.

### 2.2 Optimization of the environment embedding

Next, we alternate to optimize ***α*** for fixed variance components ***σ***^2^, such that the learned embedding ***e*** = ***Eα*** concentrates G × E signal while accounting for N × E heteroskedasticity. In particular, we select a single environmental embedding such that the induced kernel ***K***_*g*×*e*_(***α***) aligns with phenotype covariance while penalizing variance inflation captured by ***K***_*n*×*e*_(***α***). For notational simplicity, we write ***K***_*g*×*e*_ = ***K***_*g*×*e*_(***α***), ***K***_*n*×*e*_ = ***K***_*n*×*e*_(***α***). Based on the objective in Eq. (2), the terms depending on ***α*** simplify to (see Notes S1):

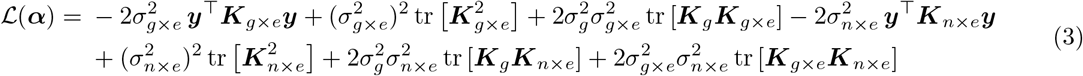

The Euclidean gradient ∇_***α***_ ℒ(***α***) is then derived as:

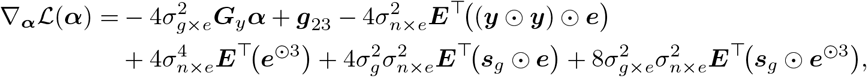

where

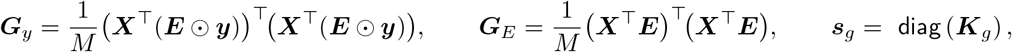

and ***e***^⊙3^ is the element-wise cube of ***e*** (Table S1). The vector ***g***_23_ aggregates the trace-gradient contributions from the second and third terms; see Section 2.3 for efficient evaluation.

Note that there is a scale non-identifiability between ***α*** and 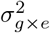: scaling ***α*** rescales ***K***_*g*×*e*_ and can be offset by 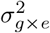. We therefore fix ‖***α*** ‖ _2_ = 1 and optimize on the unit sphere 𝕊^*L*−1^. Defining 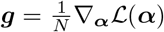, we project to the tangent space at ***α*** via ***g***_⊤_ = ***g*** − (***α*** ^⊤^***g***) ***α*** and take a capped geodesic step:

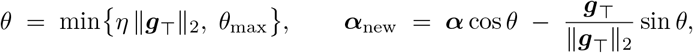

where *θ*_max_ is the maximum allowed step size. This exactly preserves unit norm and caps the per-iteration rotation, stabilizing progress when gradients are large. In practice, our proposed scheme of alternating (i) solving the 4 ×4 MoM system for 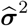 at the current ***α*** and (ii) taking one or a few geodesic steps on ***α*** converges quickly in our experiments.

### 2.3 Efficient learning via streaming and pre-computation

The primary computational challenge is that both solving for 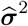 and ***α***-gradient require repeated contractions with genotype-derived kernels. Naïvely rebuilding genotype-based quantities at each iteration is prohibitively expensive at biobank scale. Instead, we perform a single streamed pass over the genotype matrix *X* to cache probe sketches that suffice for all subsequent evaluations, thereby decoupling per-iteration cost from the number of SNPs *M*.

We employ Hutchinson’s trace estimator [34], using *B* ≪ *M* independent standard-normal probe vectors collected as the rows of ***W*** ∈ ℝ^*B*×*N*^. Rather than forming the kernel matrices, we multiply each target matrix by the random probes to obtain compact sketches. Traces are approximated by the average Frobenius inner product of the corresponding sketches; accordingly, we compute kernel sketches ***U*** _*r*_ = ***K***_*r*_***W*** ^⊤^ so that the pairwise kernel trace tr [***K***_*r*_***K***_*s*_] follows from 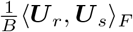. For the additive kernel, ***U*** _*g*_ is fixed and computed once. Meanwhile, the context-specific kernel ***K***_*g*×*e*_ depends on ***α*** only through ***e*** = ***Eα*** and admits a linear decomposition over environments. Let ***Z***_ℓ_= ***X*** ⊙ ***E***_:, ℓ_ be the per-environment modulated genotype matrix. During the single streamed pass over SNP blocks, we compute and cache pairwise probe sketches

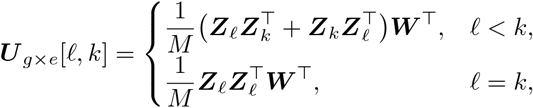

so that for any ***α*** we can assemble

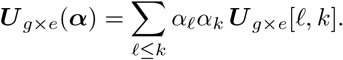

Then, both trace and cross-trace terms involving *K*_*g*×*e*_ may be computed as:

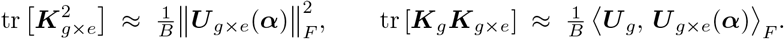

To obtain the trace-gradient ***g***_23_ without forming a *L* × *L* matrix, we first compute scalar contractions

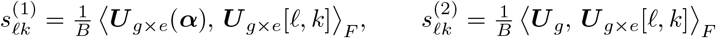

Next, we define the upper-triangular matrix ***C*** ∈ ℝ^*L*×*L*^ as:

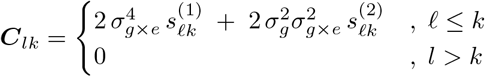

Then, we derive ***g***_23_ = (***C***+***C***^⊤^)***α***. In practice, we realize ***g***_23_ via two indexed scatter-adds over the *P* = *L*(*L*+1)*/*2 pairs [35–37], avoiding materializing ***C*** while injecting ***α*** into all trace terms at a cost that is independent of *M*.

Finally, to bound memory, we stream the genotype matrix in SNP blocks, updating all sketches and quadratic forms in-place and discarding block-level temporaries. After this single pass, subsequent iterations reuse the cached quantities: each MoM solves (the 4 ×4 system) and each ***α***-gradient evaluation runs in time independent of *M*, depending only on the number of individuals *N*, the number of environments *L*, and the number of random probes *B*. Therefore, streaming and pre-computation provide biobank-scale efficiency without sacrificing the statistical structure of the model.

### 2.4 Regularized cross-fitting for G×E

As outlined in Algorithm 1, our procedure alternates between estimating the variance components at the current environment embedding and updating the environment weights to decrease the loss in Eq. (3). While computationally convenient, reusing the same sample for both steps couples learning and evaluation and can introduce target leakage, with the environment weights adapting to noise in the phenotype vector. In the presence of a nonzero genetic main effect 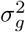 or environment-specific noise 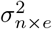, the optimizer may synthesize artificial G×E by borrowing signal from the main effect or from heteroskedasticity. To address overfitting, we introduce variant-level cross-fitting and an *ℓ*_1_-regularized update on the sphere to break this coupling and stabilize the embedding.

To decouple learning and evaluation, we use cross-fitting over variants. We split *M* SNPs into two disjoint halves (e.g., by chromosome or LD blocks, balancing MAF/LD), obtaining matrices ***X***_(1)_, ***X***_(2)_ ∈ ℝ^*N*×*M/*2^. Using only ***X***_(1)_, we make a single streamed pass to build the required sketches and optimize a unit-norm embedding ***α***_(1)_. Freezing this embedding, we then use only ***X***_(2)_ to assemble ***A***_(2)_ and ***b***_(2)_, solve for 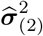, and compute standard errors. We then swap roles: learn *α*_(2)_ on ***X***_(2)_, and evaluate 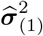 on ***X***_(1)_. This separation prevents ***α*** from capturing noise in the same variants used for component estimation. We derive the final embedding as the average of the two held-out solutions followed by renormalization to unit norm, and evaluate variance components on the full dataset to capture variance explained by all SNPs.

Cross-fitting removes the main leakage path, but dense embeddings can still overfit by combining many weak, correlated environments. To constrain model complexity and improve interpretability, we add *ℓ*_1_ regularization on the sphere, solving

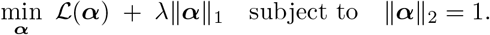

The unit-norm constraint fixes the scale ambiguity with 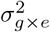, and the *ℓ*_1_ penalty encourages a sparse set of nonzero weights. This reduces effective degrees of freedom, improves robustness under collinearity, and concentrates the embedding on environments with strongest support.

We implement the update in each coordinate via soft-thresholding followed by renormalization,

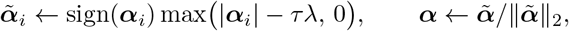

where *τ* is the step length along the geodesic move. This “soft-threshold on the sphere” integrates directly into the existing update with negligible overhead. Because a fixed, strong penalty can bias the solution, we apply light annealing: start with large *λ* = *λ*_0_ to obtain a stable coarse embedding, then decay *λ* toward a lower bound *λ*_min_ across outer alternations.

Together, these changes control overfitting. Cross-fitting separates learning ***α*** from evaluating ***σ***^2^, blocking gains that arise from relabeling noise as signal. The *ℓ*_1_ term reduces the tendency to inflate ‖ ***K***_*g*×*e*_ ‖_*F*_ or mimic ***K***_*g*_ without improving fit, and annealing allows the procedure to begin conservatively and refine as evidence accumulates. As a result, the embedding is more stable across resamples and interaction estimates are better calibrated under the null without sacrificing ENGINE’s one-pass, streamed MoM efficiency.

## 3 Results

### 3.1 Calibration and power

We benchmarked ENGINE against the most relevant alternative, LEMMA, in terms of the calibration under the null of no G × E. We simulated phenotypes for *N* = 10,000 unrelated UK Biobank (UKB) participants of White British ancestry using *M* = 10,000 SNPs genotyped on the UKB array (see Notes S4). We used five lifestyle environments (Townsend deprivation index, sleep duration, age, smoking status, alcohol frequency), mixing categorical and quantitative measures to assess robustness to different value types. Phenotypes followed Model 1 with 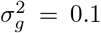 and an environmental score ***Eα***, where ***α*** ∈ ℝ^5^ was simulated from a standard normal distribution. We varied the heteroskedastic noise 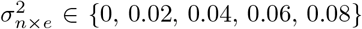 while fixing 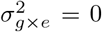, and ran 200 replicates per setting, comparing ENGINE with LEMMA. When 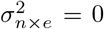 ENGINE produced 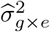 centered at zero, whereas LEMMA exhibited a slight positive bias (mean ≈ 0.010 across replicates; Figure 2A). One plausible source is data reuse: the environmental score is learned and G×E heritability is estimated on the same data, inducing overfitting (also observed for ENGINE without regularization; see Section 3.3). As 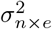 increased, ENGINE remained unbiased (overall mean across all settings − 8.8 × 10^− 6^), while the bias associated with LEMMA increased with heteroskedasticity. Type-I error was well calibrated for ENGINE: at significance threshold *p* < 0.05, the observed rejection proportions lay within their 95% binomial intervals at all 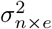 levels. In contrast, LEMMA showed inflated rejection rates both with and without heteroskedastic noise (Figure 2B).

**Figure 2.**
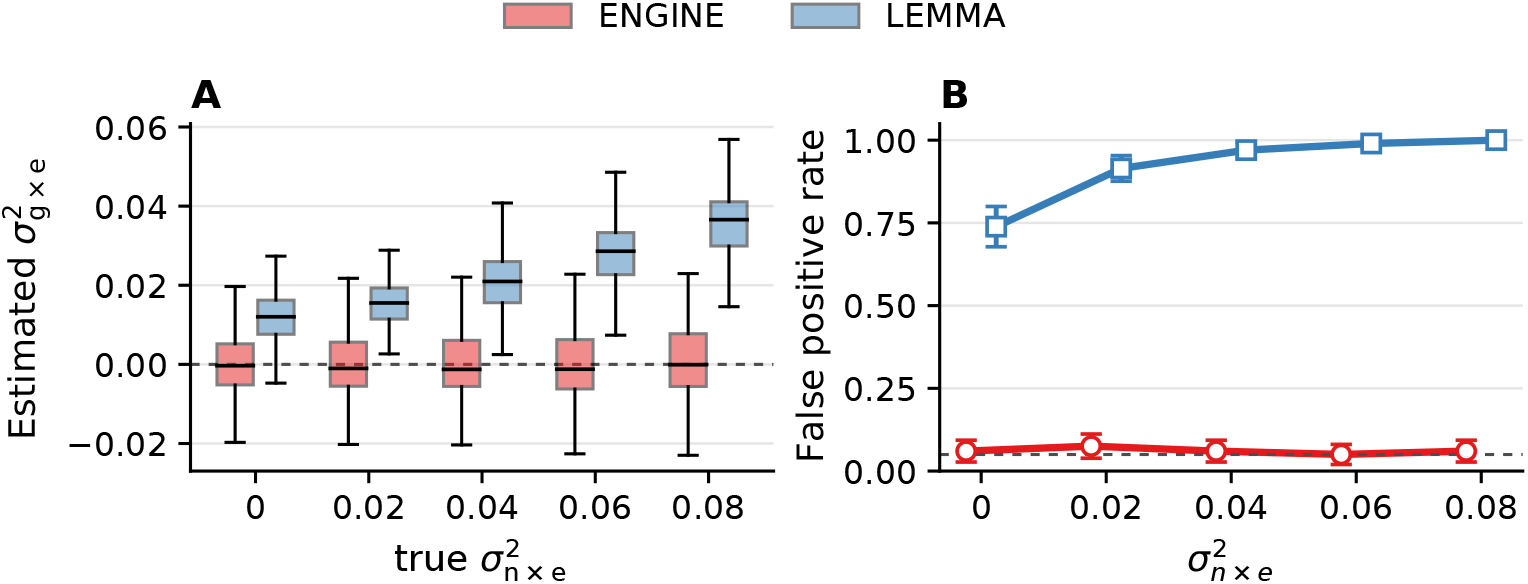
Calibration under the null across varying levels of heteroskedastic noise. We simulated phenotypes from *N* = 10,000 individuals and *M* = 10,000 SNPs with five lifestyle environments under the null of no G×E variance 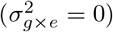, varying noise-by-environment variance 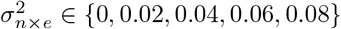 across 200 replicates. **A. Estimated** 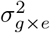. Boxplots across 200 replicates; ENGINE is marked in blue and LEMMA in orange. **B. Type I error**. False-positive rate at significance level 0.05 with 95% binomial confidence intervals.

We next evaluated the statistical power of ENGINE when G×E is present. We fixed 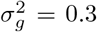, varied 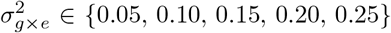, and set 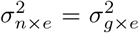,. ENGINE yielded accurate variance-component estimates: 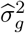 was centered on the truth across all settings, and 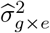 tracked the simulated values closely as 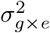 increased (Figure 3A). Power, calculated as the proportion of p < 0.05 rejections, approached 1 by 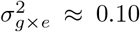 (Figure 3B). We then assessed recovery of the environment embedding using a sign-invariant metric: the squared correlation *r*^2^ between the predicted and true embeddings, which can be interpreted as the fraction of variation in the true embedding explained by the learned embedding. ENGINE achieved *r*^2^ > 0.9 when 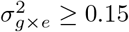 (Fig. 3C). We also confirmed in simulations that the number of random probe vectors *B* does not affect the accuracy of ENGINE in prediction (Figure S1). We used *B* = 100 by default in all experiments. Together, these results indicate that ENGINE is well calibrated under the null, powerful when interactions are present, and accurately recovers both variance components and the environment embedding.

**Figure 3.**
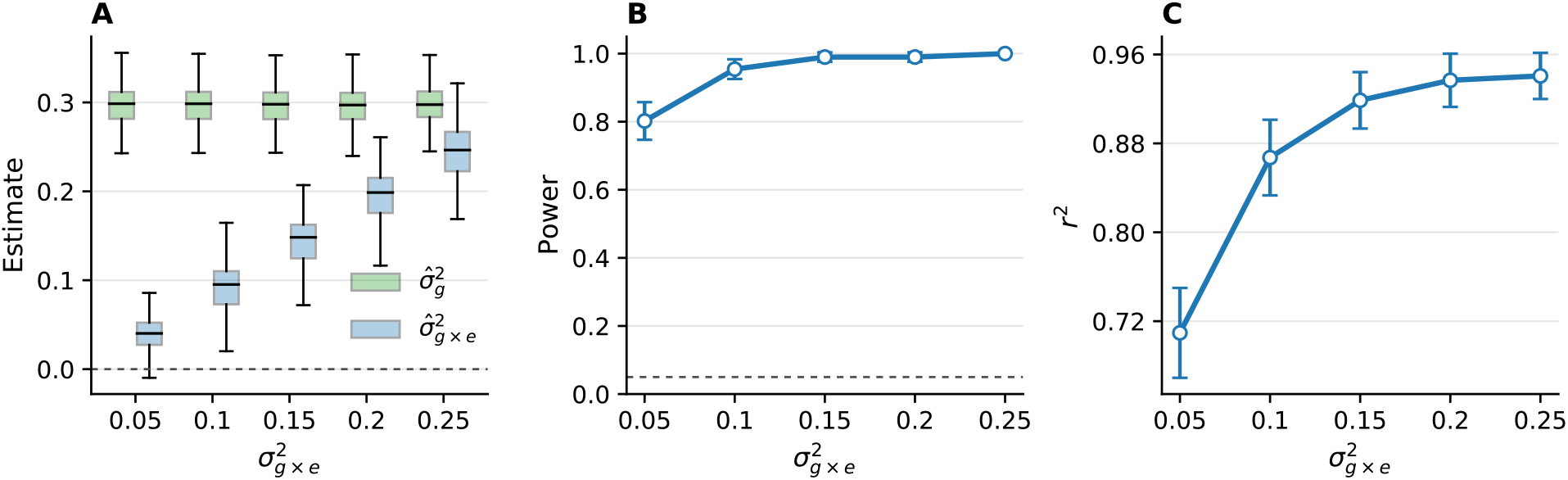
Power and estimation accuracy as a function of interaction strength. We simulated phenotypes from *N* = 10,000 individuals and *M* = 10,000 SNPs with five lifestyle environments, setting 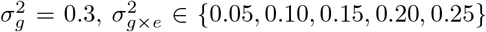, and 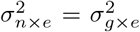 (200 replicates per setting). **A. Variance estimates**. Boxplots of 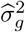 are in green and 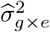 in blue across replicates; the horizontal dashed line marks zero. **B. Power**. Rejection rate at significance level 0.05 with 95% binomial confidence intervals. **C. Embedding accuracy**. Squared correlation between the learned environmental embedding and the ground truth.

### 3.3 Efficiency and scaling

We benchmarked ENGINE against LEMMA and the exact-per-update ENGINE baseline, which recomputes the exact traces and rebuilds the normal equations at every embedding update. Recall that the efficient implementation of ENGINE makes a single streaming 𝒪 (*NM*) pass over the genotype matrix to cache sketches of GRMs; subsequent embedding updates reuse these caches without scanning the raw genotypes. For the experiments below, we used five lifestyle environments by default. With SNPs fixed at *M* = 10,000, number of environmental exposures at *L* = 5 and individuals varied *N* ∈ { 1, 3, 5, 10, 30, 50, 100 } k, ENGINE is highly scalable: while LEMMA is slightly faster for smaller sample sizes when the number of individuals is less than 10, 000, its runtime grows more steeply so that for sample sizes of *N* = 100, 000 its runtime is ∼ 5 × that of ENGINE (Figure 4A). The exact-per-update ENGINE baseline is computationally prohibitive, taking ∼ 900 s at *N* = 1, 000 (∼ 45 × slower than ENGINE). In a second experiment, with individuals fixed at *N* = 10,000 and SNPs varied *M* ∈ {1, 3, 5, 10, 30, 50, 100 } k, ENGINE is marginally slower at the smallest *M* but becomes faster once the number of SNPs reaches 10, 000; the exact-per-update ENGINE baseline remains infeasible as its per-iteration cost grows with *M* (Figure 4B). In an additional experiment with both individuals and SNPs fixed at *N* = *M* = 10, 000, ENGINE exhibits similar runtime scaling in the number of environments as LEMMA (Figure S2). We further report that ENGINE fits the full UK Biobank array dataset (*M* = 454,207 variants; *N* = 291,273 individuals; *L* = 5 environments) in approximately seven hours on a single CPU core.

**Figure 4.**
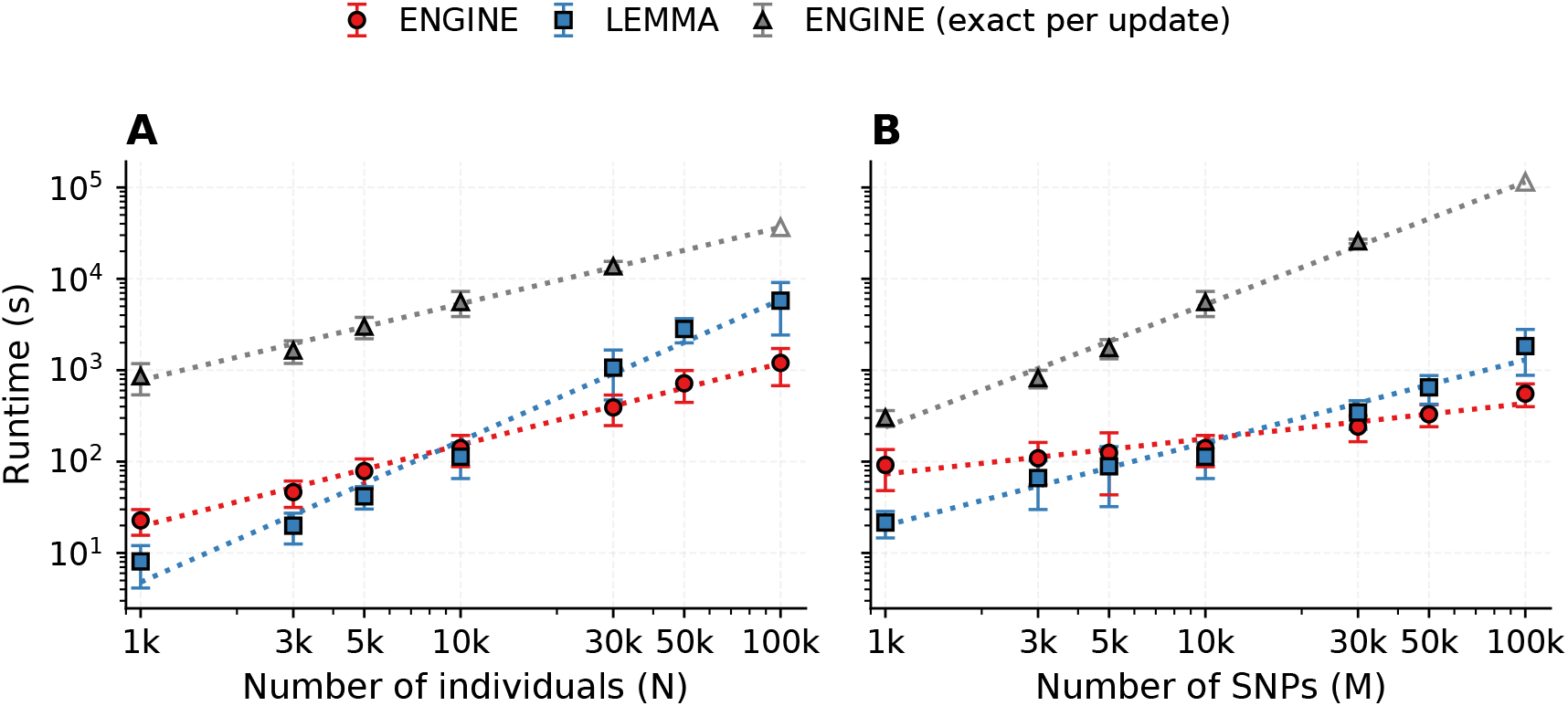
Runtime benchmarks against alternative approaches. Points show mean wall-clock runtime over 200 replicates; error bars denote one standard deviation. Dotted curves are power-law fits over the observed range, and open markers at 100 k indicate values obtained by extrapolating these fits. Both axes are on a logarithmic scale. **A. Scaling with sample size**. The number of SNPs is fixed at *M* = 10,000 and we vary the number of individuals *N* ∈ { 1, 3, 5, 10, 30, 50, 100 } k. The runtime for the exact-per-update ENGINE baseline at *N* = 100 k is extrapolated because the method did not complete within the 8-hour wall-clock limit under 30 GB of memory. **B. Scaling with SNP count**. The number of individuals is fixed at *N* = 10,000 and we vary the number of SNPs *M* ∈ { 1, 3, 5, 10, 30, 50, 100 } k. The runtime for the exact-per-update ENGINE baseline at *M* = 100 k is likewise extrapolated. All benchmarks were run on a single core of an Intel Xeon Gold 6140 CPU (36 cores, 2.30 GHz) with a maximum of 30 GB memory and an 8-hour wall-clock limit per replicate.

### 3.3 Ablation studies

We evaluated how cross-fitting (A/B holdout) and *ℓ*_1_ penalization of the environment embedding affect false-positive control under the no G×E calibration scenarios, varying heterogeneous noise 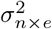 and running 100 replicates per setting at significance level 0.05; unless noted, the regularization parameter was *λ* = 5 × 10 ^− 3^. Without either safeguard, the procedure was severely inflated. At 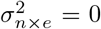, the false positive rate (FPR) was 0.48. As noise heterogeneity increased, inflation lessened but remained substantial (FPR = 0.22 at 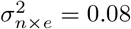). Each safeguard alone reduced inflation: cross-fitting was most effective when 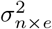 was larger, with FPR decreasing from 0.11 to 0.06 as 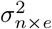 rose from 0.02 to 0.08, whereas *ℓ*_1_ regularization helped most when noise heterogeneity was absent or small (FPR ≈ 0.09 at 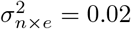) and was slightly less protective as 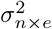 grew (rising to ≈ 0.11 at 0.08). Used together, the two mechanisms restored near-nominal behavior, with FPR ≈ 0.05 at 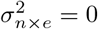 and mean FPR across settings 0.06 (95% binomial CI [0.01, 0.11]) (Figure 5). These results indicate that cross-fitting and *ℓ*_1_ regularization together enable calibrated false positive rates across the range of tested noise heteroskedasticity levels.

**Figure 5.**
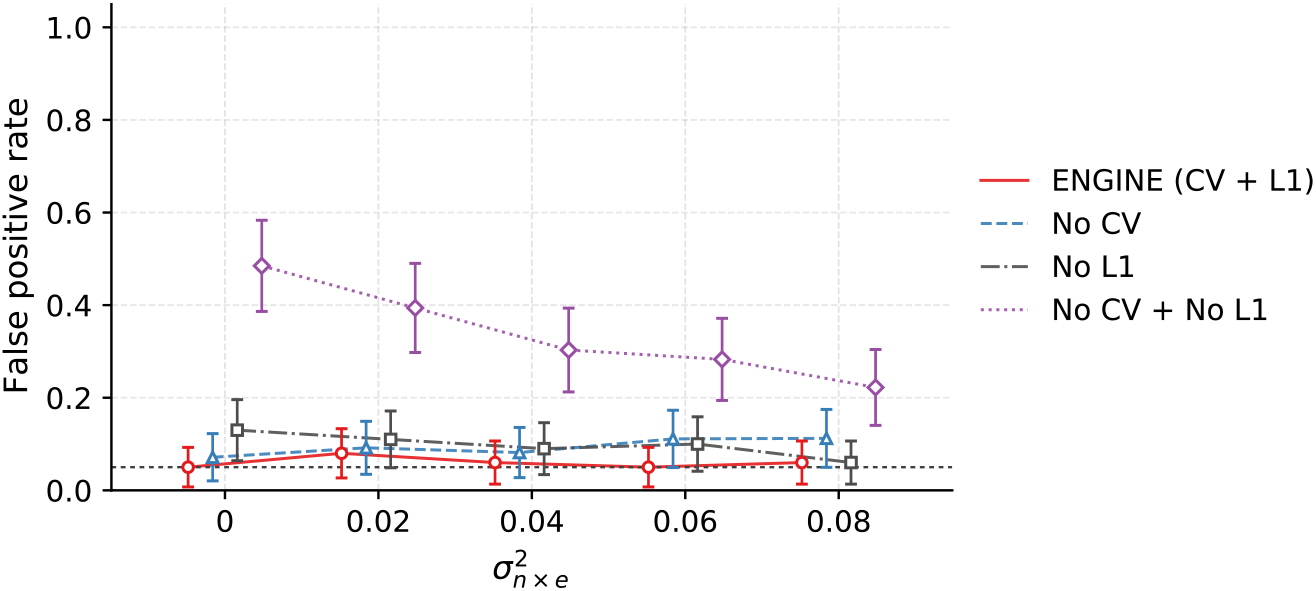
Ablation of cross-fitting (CV) and *ℓ*_1_ regularization on false-positive control. False positive rate under no true G×E as a function of heterogeneous noise 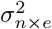. Curves compare ENGINE with both safeguards (CV+*ℓ*_1_), without CV, without *ℓ*_1_, and with neither. The false postive rate is calculated as P(rejection at *p* < 0.05) across 100 replicates; error bars show 95% binomial CIs; the dashed line marks the significance level 0.05 (*ℓ*_1_ penalty *λ* = 5 × 10^−3^).

### 3.4 G×E associated with lifestyle factors in the UK Biobank

Lifestyle context can modulate the genetic architecture of complex traits, with prior studies reporting interactions across socioeconomic status, physical activity, sleep, smoking, alcohol intake, and diet for body mass index (BMI) [38–45]. From 42 derived environments including dietary and non-dietary measures and their interactions with sex and age (Notes S3; Table S2), we pre-screened and retained ten that each showed significant 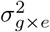 with BMI in UK Biobank, after adjusting for sex, age, and the top 20 genetic principal components. Applying ENGINE to these ten environments uncovered a substantial, interpretable lifestyle G × E component (Table S3). Using common array-genotyped SNPs with MAF > 0.01 (*M* = 454,207), we trained the environmental embedding on *N* = 50,000 individuals and then estimated variance components on the full set of *N* = 291,273 unrelated White British participants. The learned embedding yielded 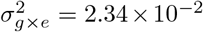 (SE: 4.2 × 10^−3^), exceeding the interaction variance attributable to any single exposure (Figure 6A).

**Figure 6.**
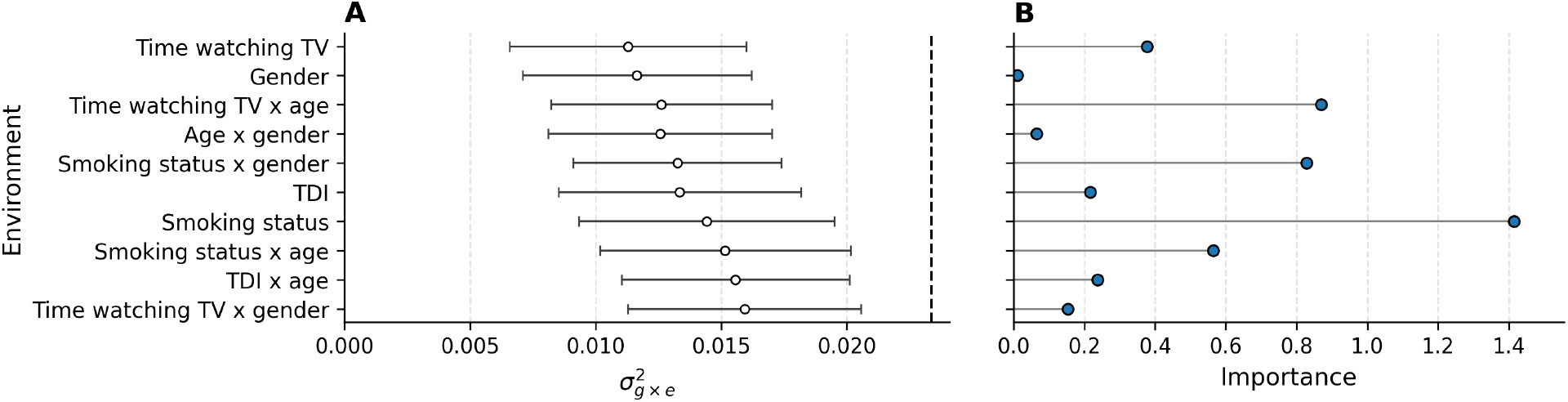
Lifestyle G×E for BMI: single environments vs. learned embedding. A. Single-exposure estimates. Estimates of 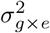 for each selected environmental exposure, with error bars indicating the estimated standard errors; the dashed vertical line marks the learned embedding estimate 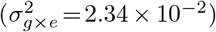). **B. Learned embedding weights**. The magnitude of learned embedding weights for the same features.

We then compared the learned embedding to unsupervised baselines constructed as the top principal component (PC) from (i) the selected ten exposures and (ii) the full set of 42 derived environments. ENGINE produced both a larger 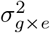 and stronger statistical evidence than either PC baseline: the estimate from the learned embedding was around 4× that of the top PC from the ten exposures 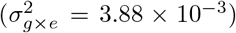 and substantially larger than the top PC from all 42 environments (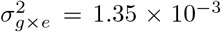 Figure 7A). A paired comparison over leave-one-out jackknife blocks (100 blocks) confirmed that blockwise estimates of 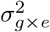 were consistently higher for the learned embedding than for the top PC of the ten exposures (one-sided *p* = 6.62 × 10^−15^; see Notes S2). Targeting the interaction structure directly thus yields measurable gains over unsupervised projections that need not align with the G×E signal. Finally, the learned embedding also exceeded the strongest single-environment signal, with 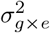 significantly larger than the maximum across individual environments (jackknife *p* = 2.67 10^−6^).

**Figure 7.**
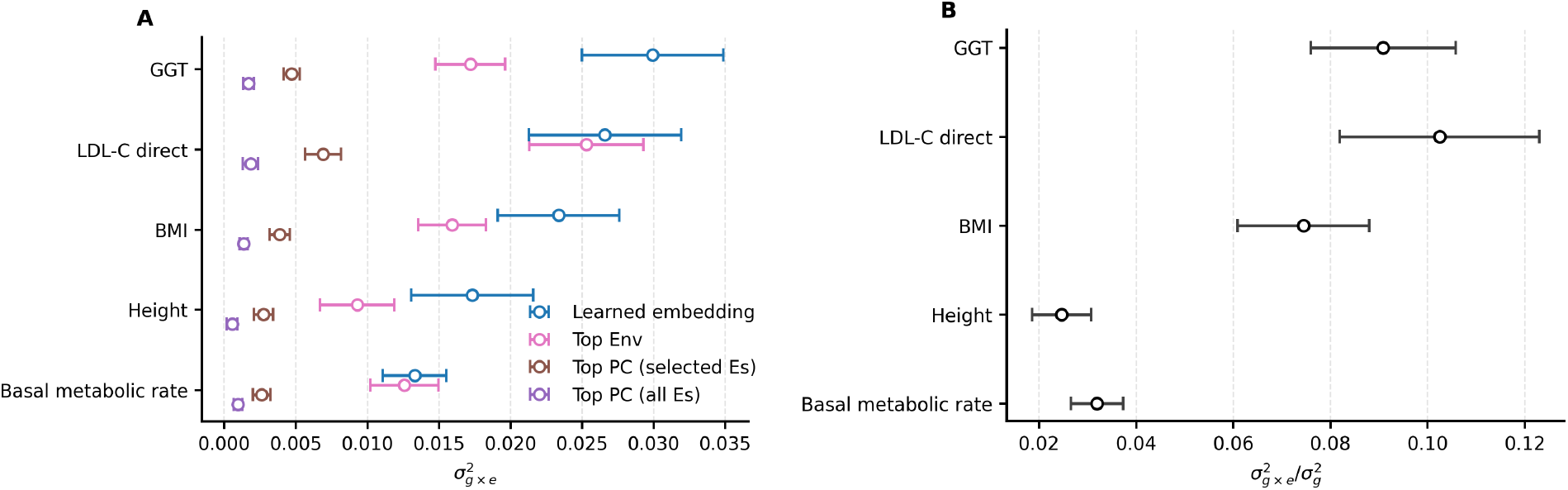
Lifestyle G ×E across traits. A. Learned embedding vs. baselines. Estimates of 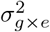 across traits (basal metabolic rate, height, BMI, LDL-C direct, GGT) for four approaches: the learned environmental embedding (blue), the best single environment (“Top Env”, pink), the top principal component computed from the selected environments (brown), and the top principal component from all environments (purple). Points show estimates; horizontal bars denote the standard errors. **B. G**×**E proportion of additive variance**. Estimates of the ratio 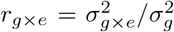 for the learned embedding across traits, with error bars indicating the estimated standard errors.

The learned weights 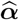 provide additional interpretability. Smoking-related measures (and their interactions) received the largest magnitudes, while nontrivial weight was also assigned to other features (e.g., time spent watching TV and TDI-related measures), supporting a diffuse architecture in which many exposures contribute modestly to the overall G ×E component (Figure 6B). This pattern aligns with prior evidence that lifestyle factors collectively shape BMI’s genetic effects [17, 32].

In addition to BMI, we extended our lifestyle-exposure analysis to four traits—basal metabolic rate, standing height, LDL cholesterol (LDL-C) direct, and gamma glutamyltransferase (GGT). For all five traits, after residualizing each trait on fixed-effect covariates (basal metabolic rate additionally adjusted for BMI), the learned embedding yielded a significantly larger estimate of 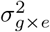 than the top PC of the same environments (jackknife *p* < 0.05*/*(42 × 5), Bonferroni-corrected across exposures and traits). To quantify the magnitude of G×E relative to additive effects, we computed 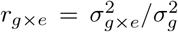 on the full dataset; across traits, this ratio ranged from 2.47% for height to 10.25% for LDL-C levels (Figure 7B). Within the context of lifestyle G ×E, these results indicate that height has a largely additive genetic architecture, whereas lifestyle exposures substantially modulate the genetic architecture of LDL-C levels [8, 31, 46–48]. Comparisons with the single environment that attains the largest 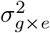 estimate show that the learned embedding captures additional G ×E signal for height and GGT, while marginally for LDL-C direct and basal metabolic rate (Figure 7A; Figure S3). These trait-specific differences indicate that not all traits exhibit substantial additional G×E beyond the contribution of the single most interacting environment.

## 4 Discussion

We present ENGINE, a scalable multi-environment G ×E estimator that learns a one-dimensional environmental embedding to concentrate interaction signal while explicitly modeling context-dependent noise. The central computational idea is to precompute genotype-dependent summaries and cache them, so each iteration can form gradients and update variance components without repeated passes over the full genotype matrix. This design makes training practical at biobank scale. Methodologically, cross-fitting and *ℓ*_1_ regularization reduce leakage from additive effects and noise-by-environment components, improving calibration while retaining strong power under heterogeneous noise.

Interpreting the learned embedding requires care. Because the interaction term is bilinear, optimization over the environment weights is nonconvex and may converge to different local optima depending on initialization. Nevertheless, we found that ENGINE yielded generally consistent embeddings and variance-component estimates across random seeds (Figure S4). In practice, stability can be improved by imposing mild structure—for example, constraining weights to be nonnegative after preprocessing and using multiple random starts to report a consensus embedding when feasible. We also note that the embedding is trained in the European-ancestry cohort; transfer to other populations depends on the degree to which causal variants and G ×E architectures are shared across ancestries. Finally, we estimate variance components via the method of moments. While potentially less statistically efficient than restricted maximum likelihood (REML) under particular assumptions [49], the method-of-moments approach is substantially faster and integrates naturally with cached summaries [50, 51].

Several directions could broaden applicability and robustness. First, cache size currently scales as *O*(*L*^2^) in the number of exposures *L*, which may be memory-limiting for hundreds to thousands of environments. A practical remedy is to compress storage using anchored sketches—for example, maintaining a single “anchor” environment sketch and storing per-exposure projections or scaling factors relative to this anchor—thereby preserving fast iterations while reducing memory. Second, correlated exposures remain challenging: we currently standardize and whiten inputs for stability, but learning directly on correlated features would improve usability and interpretability. Covariance-aware parameterizations or structured regularization that explicitly models correlation could replace preprocessing while preserving identifiability. Third, ENGINE currently learns linear combinations of exposures; incorporating modest nonlinearity (e.g., low-degree feature maps or kernelized expansions with sketching) could capture higher-order environmental effects without compromising scalability. Beyond variance decomposition, the learned embedding can be fixed and reused in downstream analyses to estimate stabilized SNP-by-embedding interaction effects, facilitating locus prioritization and biological interpretation. An additional theoretical direction is to establish convergence guarantees for sketch-based updates and error bounds that quantify approximation accuracy as a function of the number of random probe vectors.

Overall, ENGINE provides an efficient and well-calibrated framework for discovering environmental contexts that modulate genetic effects on complex traits, bridging statistical interpretability with biobank-scale computation.

## 5 Code availability

The ENGINE software is publicly available at https://github.com/sriramlab/ENGINE.

## Acknowledgements

This research was conducted using the UK Biobank Resource under application 33127. We thank the participants of UK Biobank for making this work possible. This work was funded, in part, by NIH grants R35GM153406 (Z.L., M.J., A.A., and S.S.), 1R21HG013393 (A.R.) and NSF grant CAREER-1943497 (S.S.).

### Algorithm 1 Alternating optimization for learning the environment embedding

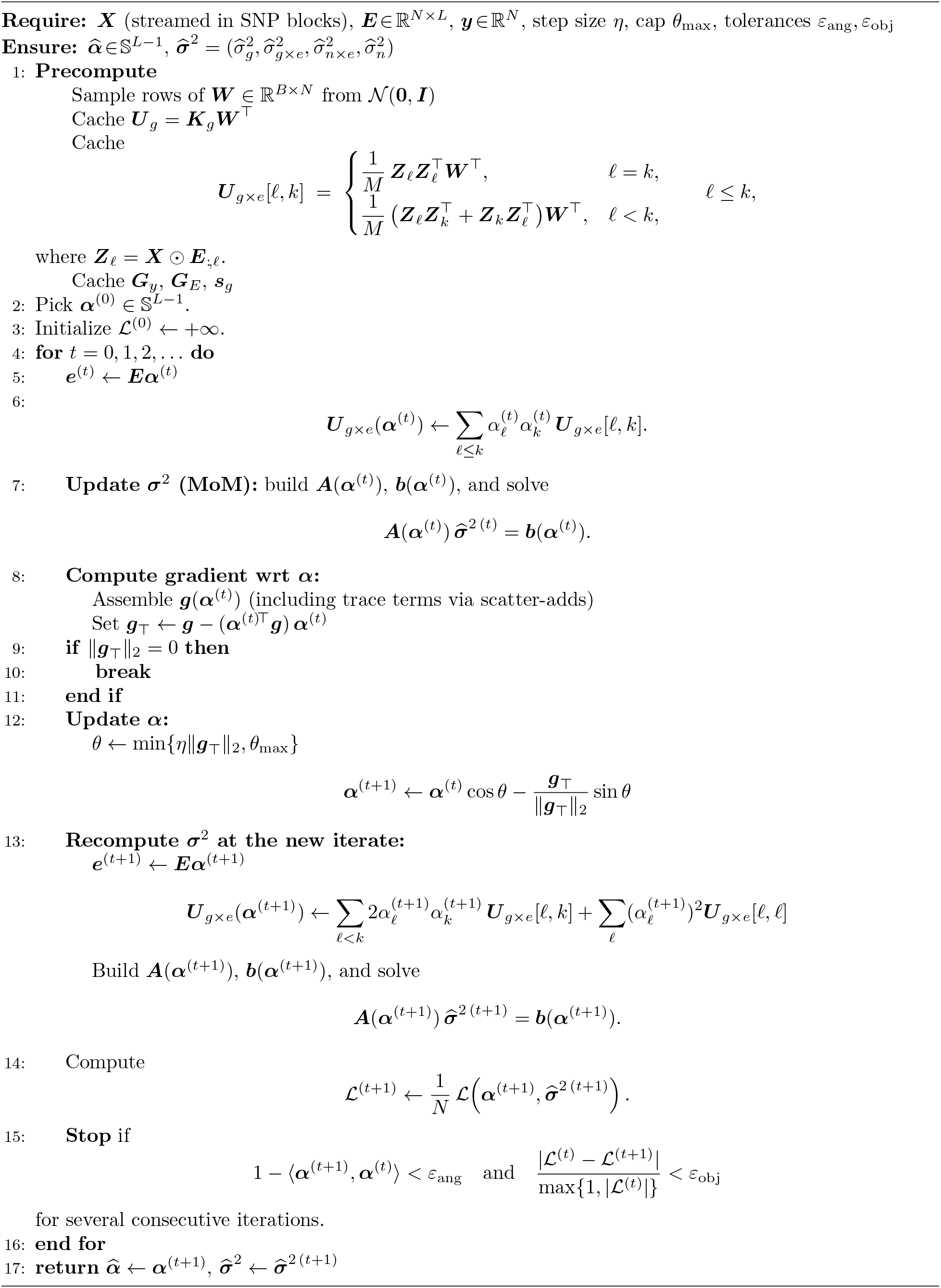

## References

[1] Esther Herrera-Luis, Kelly Benke, Heather Volk, Christine Ladd-Acosta, and Genevieve L Wojcik. Gene–environment interactions in human health. Nature Reviews Genetics, 25(11):768–784, 2024.

[2] Kenneth E Westerman and Tamar Sofer. Many roads to a gene-environment interaction. The American Journal of Human Genetics, 111(4):626–635, 2024.

[3] Samuel J Virolainen, Andrew VonHandorf, Kenyatta CMF Viel, Matthew T Weirauch, and Leah C Kottyan. Gene–environment interactions and their impact on human health. Genes & Immunity, 24 (1):1–11, 2023.

[4] Carly Boye, Shreya Nirmalan, Ali Ranjbaran, and Francesca Luca. Genotype environment interactions in gene regulation and complex traits. Nature Genetics, 56(6):1057–1068, 2024.

[5] Shinichi Namba, Kyuto Sonehara, Yuriko N Koyanagi, Takezo Kikuchi, Takafumi Ojima, Ryuya Edahiro, Go Sato, Taiki Yamaji, Yoshihiko Tomofuji, Hiroyuki Ueda, et al. A cross-population compendium of gene–environment interactions. Nature, pages 1–10, 2026.

[6] Eric Weine, Samuel Pattillo Smith, Rebecca Kathryn Knowlton, and Arbel Harpak. Trade-offs in modeling context dependency in complex trait genetics. Elife, 13:RP99210, 2025.

[7] Andy Dahl, Khiem Nguyen, Na Cai, Michael J Gandal, Jonathan Flint, and Noah Zaitlen. A robust method uncovers significant context-specific heritability in diverse complex traits. The American Journal of Human Genetics, 106(1):71–91, 2020.

[8] Ali Pazokitoroudi, Zhengtong Liu, Andrew Dahl, Noah Zaitlen, Saharon Rosset, and Sriram Sankararaman. A scalable and robust variance components method reveals insights into the architecture of gene-environment interactions underlying complex traits. The American Journal of Human Genetics, 111(7):1462–1480, 2024.

[9] Matteo Di Scipio, Mohammad Khan, Shihong Mao, Michael Chong, Conor Judge, Nazia Pathan, Nicolas Perrot, Walter Nelson, Ricky Lali, Shuang Di, et al. A versatile, fast and unbiased method for estimation of gene-by-environment interaction effects on biobank-scale datasets. Nature Communications, 14(1): 5196, 2023.

[10] Alison A Motsinger-Reif, David M Reif, Farida S Akhtari, John S House, C Ryan Campbell, Kyle P Messier, David C Fargo, Tiffany A Bowen, Srikanth S Nadadur, Charles P Schmitt, et al. Gene-environment interactions within a precision environmental health framework. Cell Genomics, 4(7), 2024.

[11] Jingjing Li, Xiao Li, Sai Zhang, and Michael Snyder. Gene-environment interaction in the era of precision medicine. Cell, 177(1):38–44, 2019.

[12] William R Reay, Joshua R Atkins, Vaughan J Carr, Melissa J Green, and Murray J Cairns. Pharmacological enrichment of polygenic risk for precision medicine in complex disorders. Scientific reports, 10 (1):879, 2020.

[13] Christopher Paul Wild. The exposome: from concept to utility. International journal of epidemiology, 41(1):24–32, 2012.

[14] Chirag J Patel, Jayanta Bhattacharya, and Atul J Butte. An environment-wide association study (ewas) on type 2 diabetes mellitus. PloS one, 5(5):e10746, 2010.

[15] Ghassan B Hamra and Jessie P Buckley. Environmental exposure mixtures: questions and methods to address them. Current epidemiology reports, 5(2):160–165, 2018.

[16] Jonathan Sulc, Ninon Mounier, Felix Günther, Thomas Winkler, Andrew R Wood, Timothy M Frayling, Iris M Heid, Matthew R Robinson, and Zoltán Kutalik. Quantification of the overall contribution of gene-environment interaction for obesity-related traits. Nature communications, 11(1):1385, 2020.

[17] Rachel Moore, Francesco Paolo Casale, Marc Jan Bonder, Danilo Horta, Lude Franke, Inês Barroso, and Oliver Stegle. A linear mixed-model approach to study multivariate gene–environment interactions. Nature genetics, 51(1):180–186, 2019.

[18] Cathie Sudlow, John Gallacher, Naomi Allen, Valerie Beral, Paul Burton, John Danesh, Paul Downey, Paul Elliott, Jane Green, Martin Landray, et al. Uk biobank: an open access resource for identifying the causes of a wide range of complex diseases of middle and old age. PLoS medicine, 12(3):e1001779, 2015.

[19] Clare Bycroft, Colin Freeman, Desislava Petkova, Gavin Band, Lloyd T Elliott, Kevin Sharp, Allan Motyer, Damjan Vukcevic, Olivier Delaneau, Jared O’Connell, et al. The uk biobank resource with deep phenotyping and genomic data. Nature, 562(7726):203–209, 2018.

[20] All of Us Research Program Investigators. The “all of us” research program. New England Journal of Medicine, 381(7):668–676, 2019.

[21] Christopher Paul Wild. Complementing the genome with an “exposome”: the outstanding challenge of environmental exposure measurement in molecular epidemiology. Cancer Epidemiology Biomarkers & Prevention, 14(8):1847–1850, 2005.

[22] Konstantinos C Makris, Andrea Baccarelli, Edwin K Silverman, and Robert O Wright. How exposomic tools complement and enrich genomic research. Cell Genomics, 5(8), 2025.

[23] Michal Sadowski, Mike Thompson, Joel Mefford, Tanushree Haldar, Akinyemi Oni-Orisan, Richard Border, Ali Pazokitoroudi, Na Cai, Julien F Ayroles, Sriram Sankararaman, et al. Characterizing the genetic architecture of drug response using gene-context interaction methods. Cell Genomics, 4(12), 2024.

[24] Jaana A Hartiala, James R Hilser, Subarna Biswas, Aldons J Lusis, and Hooman Allayee. Gene-environment interactions for cardiovascular disease. Current atherosclerosis reports, 23(12):75, 2021.

[25] Dmitry Shungin, Wei Q Deng, Tibor V Varga, Jian’an Luan, Evelin Mihailov, Andres Metspalu, GIANT Consortium, Andrew P Morris, Nita G Forouhi, Cecilia Lindgren, et al. Ranking and characterization of established bmi and lipid associated loci as candidates for gene-environment interactions. PLoS genetics, 13(6):e1006812, 2017.

[26] Guiyan Ni, Julius Van Der Werf, Xuan Zhou, Elina Hyppönen, Naomi R Wray, and S Hong Lee. Genotype–covariate correlation and interaction disentangled by a whole-genome multivariate reaction norm model. Nature communications, 10(1):2239, 2019.

[27] Wujuan Zhong, Aparna Chhibber, Lan Luo, Devan V Mehrotra, and Judong Shen. A fast and powerful linear mixed model approach for genotype-environment interaction tests in large-scale gwas. Briefings in Bioinformatics, 24(1):bbac547, 2023.

[28] Ali Pazokitoroudi, Yue Wu, Kathryn S Burch, Kangcheng Hou, Aaron Zhou, Bogdan Pasaniuc, and Sriram Sankararaman. E”cient variance components analysis across millions of genomes. Nature communications, 11(1):4020, 2020.

[29] Yue Wu and Sriram Sankararaman. A scalable estimator of snp heritability for biobank-scale data. Bioinformatics, 34(13):i187–i194, 2018.

[30] Jian Yang, S Hong Lee, Michael E Goddard, and Peter M Visscher. Gcta: a tool for genome-wide complex trait analysis. The American journal of human genetics, 88(1):76–82, 2011.

[31] Matthew R Robinson, Geoffrey English, Gerhard Moser, Luke R Lloyd-Jones, Marcus A Triplett, Zhihong Zhu, Ilja M Nolte, Jana V van Vliet-Ostaptchouk, Harold Snieder, LifeLines Cohort Study, et al. Genotype–covariate interaction effects and the heritability of adult body mass index. Nature genetics, 49(8):1174–1181, 2017.

[32] Matthew Kerin and Jonathan Marchini. Inferring gene-by-environment interactions with a bayesian whole-genome regression model. The American Journal of Human Genetics, 107(4):698–713, 2020.

[33] Moonseong Jeong, Ali Pazokitoroudi, Zhengtong Liu, and Sriram Sankararaman. Scalable summarystatistics-based heritability estimation method with individual genotype level accuracy. Genome Research, 34(9):1286–1293, 2024.

[34] Michael F Hutchinson. A stochastic estimator of the trace of the influence matrix for laplacian smoothing splines. Communications in Statistics-Simulation and Computation, 18(3):1059–1076, 1989.

[35] M Jon Turner, Ray W Clough, Harold C Martin, and LJ Topp. Stiffness and deflection analysis of complex structures. journal of the Aeronautical Sciences, 23(9):805–823, 1956.

[36] Fred G Gustavson. Two fast algorithms for sparse matrices: Multiplication and permuted transposition. ACM Transactions on Mathematical Software (TOMS), 4(3):250–269, 1978.

[37] Iain S Duff, Michael A Heroux, and Roldan Pozo. An overview of the sparse basic linear algebra subprograms: The new standard from the blas technical forum. ACM Transactions on Mathematical Software (TOMS), 28(2):239–267, 2002.

[38] Jessica Tyrrell, Andrew R Wood, Ryan M Ames, Hanieh Yaghootkar, Robin N Beaumont, Samuel E Jones, Marcus A Tuke, Katherine S Ruth, Rachel M Freathy, George Davey Smith, et al. Gene– obesogenic environment interactions in the uk biobank study. International journal of epidemiology, 46 (2):559–575, 2017.

[39] Tuomas O Kilpeläinen, Lu Qi, Soren Brage, Stephen J Sharp, Emily Sonestedt, Ellen Demerath, Tariq Ahmad, Samia Mora, Marika Kaakinen, Camilla Helene Sandholt, et al. Physical activity attenuates the influence of fto variants on obesity risk: a meta-analysis of 218,166 adults and 19,268 children. PLoS medicine, 8(11):e1001116, 2011.

[40] Qibin Qi, Audrey Y Chu, Jae H Kang, Majken K Jensen, Gary C Curhan, Louis R Pasquale, Paul M Ridker, David J Hunter, Walter C Willett, Eric B Rimm, et al. Sugar-sweetened beverages and genetic risk of obesity. New England Journal of Medicine, 367(15):1387–1396, 2012.

[41] Nathaniel F Watson, Kathryn Paige Harden, Dedra Buchwald, Michael V Vitiello, Allan I Pack, David S Weigle, and Jack Goldberg. Sleep duration and body mass index in twins: a gene-environment interaction. Sleep, 35(5):597–603, 2012.

[42] Anne E Justice, Thomas W Winkler, Mary F Feitosa, Misa Graff, Virginia A Fisher, Kristin Young, Llilda Barata, Xuan Deng, Jacek Czajkowski, David Hadley, et al. Genome-wide meta-analysis of 241,258 adults accounting for smoking behaviour identifies novel loci for obesity traits. Nature communications, 8(1):14977, 2017.

[43] Tiange Wang, Yoriko Heianza, Dianjianyi Sun, Tao Huang, Wenjie Ma, Eric B Rimm, JoAnn E Manson, Frank B Hu, Walter C Willett, and Lu Qi. Improving adherence to healthy dietary patterns, genetic risk, and long term weight gain: gene-diet interaction analysis in two prospective cohort studies. bmj, 360, 2018.

[44] Ming Ding, Christina Ellervik, Tao Huang, Majken K Jensen, Gary C Curhan, Louis R Pasquale, Jae H Kang, Janey L Wiggs, David J Hunter, Walter C Willett, et al. Diet quality and genetic association with body mass index: results from 3 observational studies. The American Journal of Clinical Nutrition, 108 (6):1291–1300, 2018.

[45] Matthew R Robinson, Gibran Hemani, Carolina Medina-Gomez, Massimo Mezzavilla, Tonu Esko, Konstantin Shakhbazov, Joseph E Powell, Anna Vinkhuyzen, Sonja I Berndt, Stefan Gustafsson, et al. Population genetic differentiation of height and body mass index across europe. Nature genetics, 47 (11):1357–1362, 2015.

[46] Loïc Yengo, Sailaja Vedantam, Eirini Marouli, Julia Sidorenko, Eric Bartell, Saori Sakaue, Marielisa Graff, Anders U Eliasen, Yunxuan Jiang, Sridharan Raghavan, et al. A saturated map of common genetic variants associated with human height. Nature, 610(7933):704–712, 2022.

[47] Aline Jelenkovic, Yoon-Mi Hur, Reijo Sund, Yoshie Yokoyama, Sisira H Siribaddana, Matthew Hotopf, Athula Sumathipala, Fruhling Rijsdijk, Qihua Tan, Dongfeng Zhang, et al. Genetic and environmental influences on adult human height across birth cohorts from 1886 to 1994. Elife, 5:e20320, 2016.

[48] Vincent Laville, Timothy Majarian, Yun J Sung, Karen Schwander, Mary F Feitosa, Daniel I Chasman, Amy R Bentley, Charles N Rotimi, L Adrienne Cupples, Paul S de Vries, et al. Gene-lifestyle interactions in the genomics of human complex traits. European Journal of Human Genetics, 30(6):730–739, 2022.

[49] H Desmond Patterson and Robin Thompson. Recovery of inter-block information when block sizes are unequal. Biometrika, 58(3):545–554, 1971.

[50] Xiang Zhou. A unified framework for variance component estimation with summary statistics in genome-wide association studies. The annals of applied statistics, 11(4):2027, 2017.

[51] Kun Yue, Jing Ma, Timothy Thornton, and Ali Shojaie. Rehe: Fast variance components estimation for linear mixed models. Genetic epidemiology, 45(8):891–905, 2021.

