## Supplementary Materials for "A biobank-scale method for learning modulators of gene-environment interaction underlying human complex traits from multiple environmental exposures"

### Supplemental Notes

#### S1 Derivation of the objective for optimization of $\boldsymbol{\alpha}$

To learn an embedding $\boldsymbol{\alpha}$ such that the variance of the phenotype is explained by the additive, G×E, and N×E components, we consider the MoM objective

$$\mathcal{L}(\boldsymbol{\alpha}, \boldsymbol{\sigma}^2) = \left\| \boldsymbol{y}\boldsymbol{y}^\top - \sigma_g^2 \boldsymbol{K}_g - \sigma_{g\times e}^2 \boldsymbol{K}_{g\times e}(\boldsymbol{\alpha}) - \sigma_{n\times e}^2 \boldsymbol{K}_{n\times e}(\boldsymbol{\alpha}) \right\|_F^2.$$

For notational convenience, define

$$\boldsymbol{A} = \boldsymbol{y}\boldsymbol{y}^\top - \sigma_g^2 \boldsymbol{K}_g, \qquad \boldsymbol{B}(\boldsymbol{\alpha}) = \sigma_{g\times e}^2 \boldsymbol{K}_{g\times e}(\boldsymbol{\alpha}) + \sigma_{n\times e}^2 \boldsymbol{K}_{n\times e}(\boldsymbol{\alpha}).$$

Then

$$\begin{aligned}
\mathcal{L}(\boldsymbol{\alpha}, \boldsymbol{\sigma}^2) &= \|\boldsymbol{A} - \boldsymbol{B}(\boldsymbol{\alpha})\|_F^2 \\
&= \|\boldsymbol{A}\|_F^2 + \|\boldsymbol{B}(\boldsymbol{\alpha})\|_F^2 - 2\,\mathrm{tr}\left[\boldsymbol{A}^\top \boldsymbol{B}(\boldsymbol{\alpha})\right].
\end{aligned}$$

The term $\|\boldsymbol{A}\|_F^2$ does not depend on $\boldsymbol{\alpha}$ and can be dropped for optimization. Writing $\boldsymbol{K}_{g\times e} = \boldsymbol{K}_{g\times e}(\boldsymbol{\alpha})$ and $\boldsymbol{K}_{n\times e} = \boldsymbol{K}_{n\times e}(\boldsymbol{\alpha})$ for brevity, we obtain

$$\begin{aligned}
\mathcal{L}(\boldsymbol{\alpha}) \propto\ & \|\sigma_{g\times e}^2 \boldsymbol{K}_{g\times e} + \sigma_{n\times e}^2 \boldsymbol{K}_{n\times e}\|_F^2 - 2\,\mathrm{tr}\left[\boldsymbol{A}^\top \left(\sigma_{g\times e}^2 \boldsymbol{K}_{g\times e} + \sigma_{n\times e}^2 \boldsymbol{K}_{n\times e}\right)\right] \\
=\ & (\sigma_{g\times e}^2)^2 \mathrm{tr}\left[\boldsymbol{K}_{g\times e}^2\right] + (\sigma_{n\times e}^2)^2 \mathrm{tr}\left[\boldsymbol{K}_{n\times e}^2\right] + 2\sigma_{g\times e}^2 \sigma_{n\times e}^2 \mathrm{tr}\left[\boldsymbol{K}_{g\times e}\boldsymbol{K}_{n\times e}\right] \\
& - 2\sigma_{g\times e}^2 \mathrm{tr}\left[\boldsymbol{A}\boldsymbol{K}_{g\times e}\right] - 2\sigma_{n\times e}^2 \mathrm{tr}\left[\boldsymbol{A}\boldsymbol{K}_{n\times e}\right].
\end{aligned}$$

Substituting $\boldsymbol{A} = \boldsymbol{y}\boldsymbol{y}^\top - \sigma_g^2 \boldsymbol{K}_g$ and using symmetry of the kernels,

$$\begin{aligned}
\mathrm{tr}\left[\boldsymbol{A}\boldsymbol{K}_{g\times e}\right] &= \mathrm{tr}\left[\boldsymbol{y}\boldsymbol{y}^\top \boldsymbol{K}_{g\times e}\right] - \sigma_g^2 \mathrm{tr}\left[\boldsymbol{K}_g \boldsymbol{K}_{g\times e}\right] = \boldsymbol{y}^\top \boldsymbol{K}_{g\times e} \boldsymbol{y} - \sigma_g^2 \mathrm{tr}\left[\boldsymbol{K}_g \boldsymbol{K}_{g\times e}\right], \\
\mathrm{tr}\left[\boldsymbol{A}\boldsymbol{K}_{n\times e}\right] &= \mathrm{tr}\left[\boldsymbol{y}\boldsymbol{y}^\top \boldsymbol{K}_{n\times e}\right] - \sigma_g^2 \mathrm{tr}\left[\boldsymbol{K}_g \boldsymbol{K}_{n\times e}\right] = \boldsymbol{y}^\top \boldsymbol{K}_{n\times e} \boldsymbol{y} - \sigma_g^2 \mathrm{tr}\left[\boldsymbol{K}_g \boldsymbol{K}_{n\times e}\right].
\end{aligned}$$

Therefore,

$$\begin{aligned}
\mathcal{L}(\boldsymbol{\alpha}) \propto\ & (\sigma_{g\times e}^2)^2 \mathrm{tr}\left[\boldsymbol{K}_{g\times e}^2\right] + (\sigma_{n\times e}^2)^2 \mathrm{tr}\left[\boldsymbol{K}_{n\times e}^2\right] + 2\sigma_{g\times e}^2 \sigma_{n\times e}^2 \mathrm{tr}\left[\boldsymbol{K}_{g\times e}\boldsymbol{K}_{n\times e}\right] \\
& - 2\sigma_{g\times e}^2 \boldsymbol{y}^\top \boldsymbol{K}_{g\times e} \boldsymbol{y} - 2\sigma_{n\times e}^2 \boldsymbol{y}^\top \boldsymbol{K}_{n\times e} \boldsymbol{y} \\
& + 2\sigma_g^2 \sigma_{g\times e}^2 \mathrm{tr}\left[\boldsymbol{K}_g \boldsymbol{K}_{g\times e}\right] + 2\sigma_g^2 \sigma_{n\times e}^2 \mathrm{tr}\left[\boldsymbol{K}_g \boldsymbol{K}_{n\times e}\right].
\end{aligned}$$

In the last step, we used the identity $\|\boldsymbol{C}\|_F^2 = \mathrm{tr}\left[\boldsymbol{C}^\top \boldsymbol{C}\right]$ and the cyclic property of the trace, $\mathrm{tr}\left[\boldsymbol{v}\boldsymbol{v}^\top \boldsymbol{A}\right] = \mathrm{tr}\left[\boldsymbol{v}^\top \boldsymbol{A}\boldsymbol{v}\right] = \boldsymbol{v}^\top \boldsymbol{A}\boldsymbol{v}$, for any vector $\boldsymbol{v}$ and matrix $\boldsymbol{A}$.

#### S2 Jackknife comparison of learned embedding vs. baseline

We use a delete-a-group jackknife to compare two estimators of the G×E variance component $\sigma^2_{g\times e}$: the learned environmental embedding from ENGINE and a baseline using the top principal component (PC1) of the same environments, based on the same partition of the data into $K$ blocks. Let $\hat{\sigma}^{2(\mathrm{ENG})}_{g\times e}$ and $\hat{\sigma}^{2(\mathrm{PC1})}_{g\times e}$ denote the full-sample estimates from ENGINE and the PC1 baseline, respectively, and let $\hat{\sigma}^{2(\mathrm{ENG})}_{g\times e,(-k)}$ and $\hat{\sigma}^{2(\mathrm{PC1})}_{g\times e,(-k)}$ denote the corresponding leave-one-block-out (LOO) estimates obtained by omitting block $k \in \{1,\dots,K\}$.

With block sizes $n_k$ and total sample size $N = \sum_{k=1}^{K} n_k$, we form delete-a-group jackknife pseudo-values

$$\tilde{\sigma}^{2(\mathrm{ENG})}_{k} = N\,\hat{\sigma}^{2(\mathrm{ENG})}_{g\times e} - \big(N - n_k\big)\,\hat{\sigma}^{2(\mathrm{ENG})}_{g\times e,(-k)}, \qquad \tilde{\sigma}^{2(\mathrm{PC1})}_{k} = N\,\hat{\sigma}^{2(\mathrm{PC1})}_{g\times e} - \big(N - n_k\big)\,\hat{\sigma}^{2(\mathrm{PC1})}_{g\times e,(-k)}.$$

We then form paired differences

$$d_k = \tilde{\sigma}^{2(\mathrm{ENG})}_{k} - \tilde{\sigma}^{2(\mathrm{PC1})}_{k}, \qquad k = 1,\dots,K,$$

with mean $\bar{d} = K^{-1}\sum_{k=1}^{K} d_k$ and sample variance

$$s_d^2 = \frac{1}{K-1}\sum_{k=1}^{K}(d_k - \bar{d})^2.$$

The jackknife standard error for the difference in $\sigma^2_{g\times e}$ is

$$\widehat{\mathrm{SE}}\big(\hat{\sigma}^{2(\mathrm{ENG})}_{g\times e} - \hat{\sigma}^{2(\mathrm{PC1})}_{g\times e}\big) = \frac{s_d}{\sqrt{K}},$$

and the $t$-statistic

$$t = \frac{\bar{d}}{s_d/\sqrt{K}}$$

is compared to a $t$ distribution with $K-1$ degrees of freedom. When testing whether the learned embedding captures strictly more G×E variance than PC1 (i.e. $H_1\colon\ \sigma^{2(\mathrm{ENG})}_{g\times e} > \sigma^{2(\mathrm{PC1})}_{g\times e}$), we report the one-sided $p$-value

$$p_{\mathrm{ENG}>\mathrm{PC1}} = 1 - F_{t_{K-1}}(t),$$

where $F_{t_{K-1}}$ is the CDF of the $t_{K-1}$ distribution.

#### S3 Data description

We analyzed UK Biobank genotype data under application 33127, including only participants with consent as verified by UKB. We restricted variants to SNPs present on the UK Biobank Axiom array and excluded SNPs with more than 1% missingness or minor allele frequency below 1%. We further removed SNPs that deviated from Hardy–Weinberg equilibrium at a significance threshold of $10^{-7}$. Our sample was limited to self-reported White British individuals with at most third-degree relatedness, defined as having all pairwise kinship coefficients $< 1/2^{(9/2)}$, and we excluded individuals who were outliers for genotype heterozygosity or missingness. After quality control, the final dataset comprised $N = 291{,}273$ individuals and $M = 454{,}207$ SNPs for real-data analysis. During training, we selected a random subset of $N = 50,000$ individuals.

For the lifestyle environmental exposures, we followed Kerin and Marchini [32] to curate a set of 42 lifestyle variables in UK Biobank. In brief, this set comprised 10 non-dietary exposures (including physical activity and sleep duration) and 11 dietary measurements (such as meat intake and alcohol consumption). We first converted raw text encodings to numerical values. For seven-level intake variables, we mapped "Never" to 0, "Less than once a week" to 1, "Once a week" to 2, "2–4 times a week" to 3, "5–6 times a week" to 4, "Once or more daily" to 5, and "Do not know" to missing. For salt added to food, we mapped "Never/rarely" to 1, "Sometimes" to 2, "Usually" to 3, and "Always" to 4. For alcohol frequency, we mapped "Daily or almost daily" to 1, "Three or four times a week" to 2, "Once or twice a week" to 3, "One to three times a month" to 4, "Special occasions only" to 5, and "Never" to 6. For smoking status, we coded "Never" as 0, "Previous" as 1, and "Current" as 2. For sex, we encoded female as 1 and male as 2.

We removed values above the 99th percentile for cooked vegetable intake and tea intake. For "Time spent watching television", values less than 0.5 hours were set to 0.5 hours, and we removed the top and bottom 1% of observations for both sleep duration and time spent watching television. For a quadratic sleep term, we computed the squared deviation from the mean sleep duration for each individual. We further augmented the non-dietary exposures with interaction terms obtained by multiplying each with sex and age. Including sex itself, this yielded a total of 42 environmental exposures, which we standardized prior to analysis. For the five focal phenotypes, we applied an inverse normal transformation. The UK Biobank fields used in our analysis are listed in Table S2. In addition, when estimating variance components on the full set of individuals, we included sex, age, and the top 20 genetic principal components as covariates.

#### S4 Simulation details

We simulated phenotypes using UK Biobank Array genotypes. Let $\boldsymbol{X} \in \mathbb{R}^{N \times M}$ denote the standardized genotype matrix. Under Eq. 1, we generated

$$\boldsymbol{y} \; = \; \boldsymbol{X}\boldsymbol{\beta} \; + \; (\boldsymbol{X} \odot \boldsymbol{e})\boldsymbol{\gamma} \; + \; (\boldsymbol{I} \odot \boldsymbol{e})\boldsymbol{\delta} \; + \; \boldsymbol{\epsilon},$$

where $\boldsymbol{\beta} \sim \mathcal{N}\big(\boldsymbol{0}, \frac{\sigma_g^2}{M}\boldsymbol{I}_M\big)$, $\boldsymbol{\gamma} \sim \mathcal{N}\big(\boldsymbol{0}, \frac{\sigma_{g\times e}^2}{M}\boldsymbol{I}_M\big)$, $\boldsymbol{\delta} \sim \mathcal{N}\big(\boldsymbol{0}, \sigma_{n\times e}^2\boldsymbol{I}_N\big)$, and $\boldsymbol{\epsilon} \sim \mathcal{N}\big(\boldsymbol{0}, \sigma_e^2\boldsymbol{I}_N\big)$. The environment score is defined as $\boldsymbol{e} = \boldsymbol{E}\boldsymbol{\alpha}$, where $\boldsymbol{\alpha} \sim \mathcal{N}(\boldsymbol{0}, \boldsymbol{I})$ is drawn and then normalized to have $\|\boldsymbol{\alpha}\|_2 = 1$. We assumed an infinitesimal polygenic architecture: for each SNP $j$, $\beta_j \sim \mathcal{N}(0, \sigma_g^2/M)$ and $\gamma_j \sim \mathcal{N}(0, \sigma_{g\times e}^2/M)$. For the NxE component, we used the identity $(\boldsymbol{I} \odot \boldsymbol{e})\boldsymbol{\delta} = \boldsymbol{e} * \boldsymbol{\delta}$, i.e., an element-wise product. Finally, we standardized $\boldsymbol{y}$ to have mean 0 and standard deviation 1.

We randomly selected $N$ = 10,000 individuals from 291,273 unrelated White British participants and used the first $M = 10,000$ SNPs (by genomic position) from a set of $M$ = 454,207 common variants with MAF $> 0.1$. We considered five lifestyle environments: categorical smoking status and alcohol frequency, and quantitative measures of Townsend deprivation index, sleep duration, and age. Each environment was standardized before analysis. In null simulations, we fixed $\sigma_{g\times e}^2 = 0$ and $\sigma_g^2 = 0.1$, and varied the heteroskedastic NxE component over $\sigma_{n\times e}^2 \in \{0, 0.02, 0.04, 0.06, 0.08\}$. For power simulations, we fixed $\sigma_g^2 = 0.3$ and varied $\sigma_{g\times e}^2 = \sigma_{n\times e}^2 \in \{0.05, 0.10, 0.15, 0.20, 0.25\}$. We also generated a new $\boldsymbol{\alpha}$ in each replicate and ran 100 replicates per setting.

### Supplemental Figures

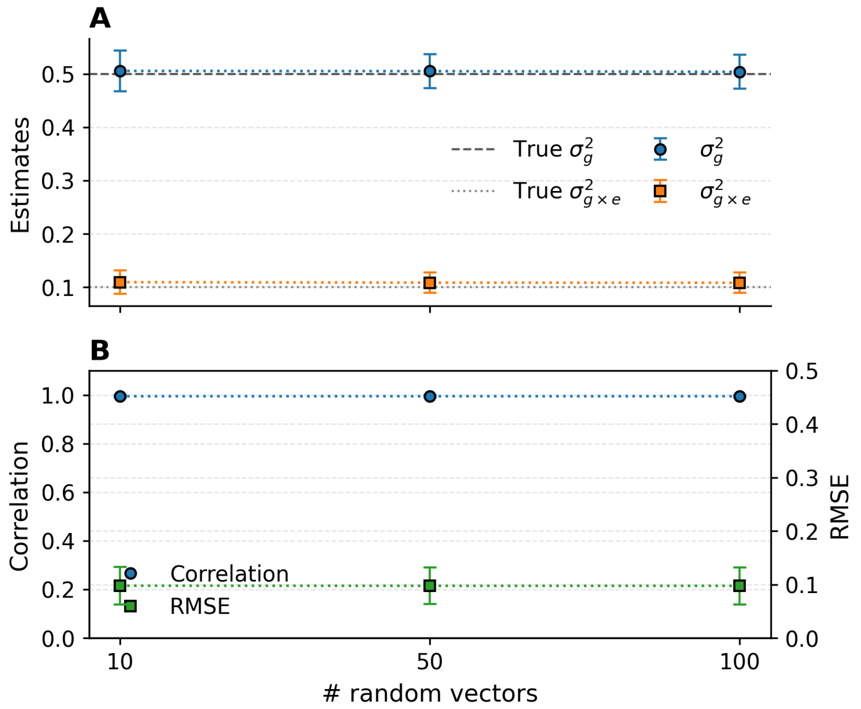


Figure S1: **Effect of the number of random vectors on ENGINE performance.** We simulated phenotypes for $N$ = 10,000 individuals and $M$ = 10,000 SNPs with $\sigma_g^2 = 0.5$ and $\sigma_{g\times e}^2 = 0.1$ across 50 replicates. Environmental scores were $\mathbf{e} = \boldsymbol{E\alpha}$ with $\boldsymbol{\alpha}$ drawn from standard normal. ENGINE was run with $B \in \{10, 50, 100\}$ random probe vectors. **A. Variance–component estimates.** Points with error bars show mean ± standard deviation for $\sigma_g^2$ (blue) and $\sigma_{g\times e}^2$ (orange); dashed and dotted lines denote the corresponding truths. **B. Recovery of the environmental embedding.** Pearson correlation (left axis, blue) and RMSE (right axis, green) between learned and true scores versus $B$. Estimates and embedding accuracy are stable across $B$.

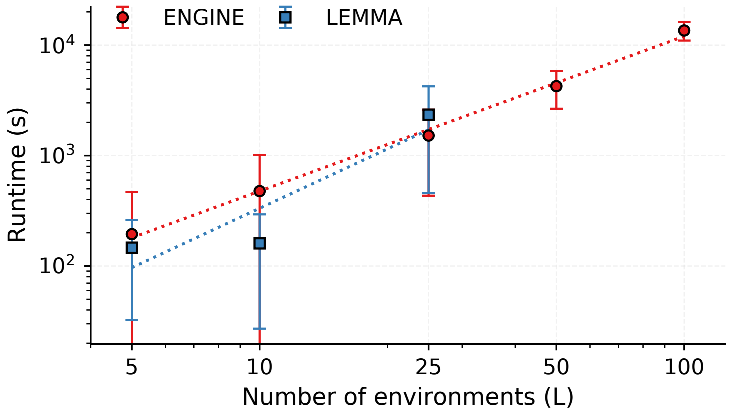


Figure S2: **Runtime benchmarks against LEMMA as a function of the number of environments.** Points show mean wall-clock runtime over 100 replicates, with error bars denoting one standard deviation. Dotted curves indicate power-law fits over the observed range. Both axes are on a logarithmic scale. We fix the sample size and number of SNPs at $N = 10{,}000$ and $M = 10{,}000$ and vary the number of environments $L \in \{5, 10, 25, 50, 100\}$. Benchmarks were run on a single core of an Intel Xeon Gold 6140 CPU (36 cores, 2.30 GHz) with at most 30 GB memory and an 8-hour wall-clock limit per replicate. LEMMA terminated with an error for $L = 50$ or $L = 100$, so the summary statistics are excluded here.

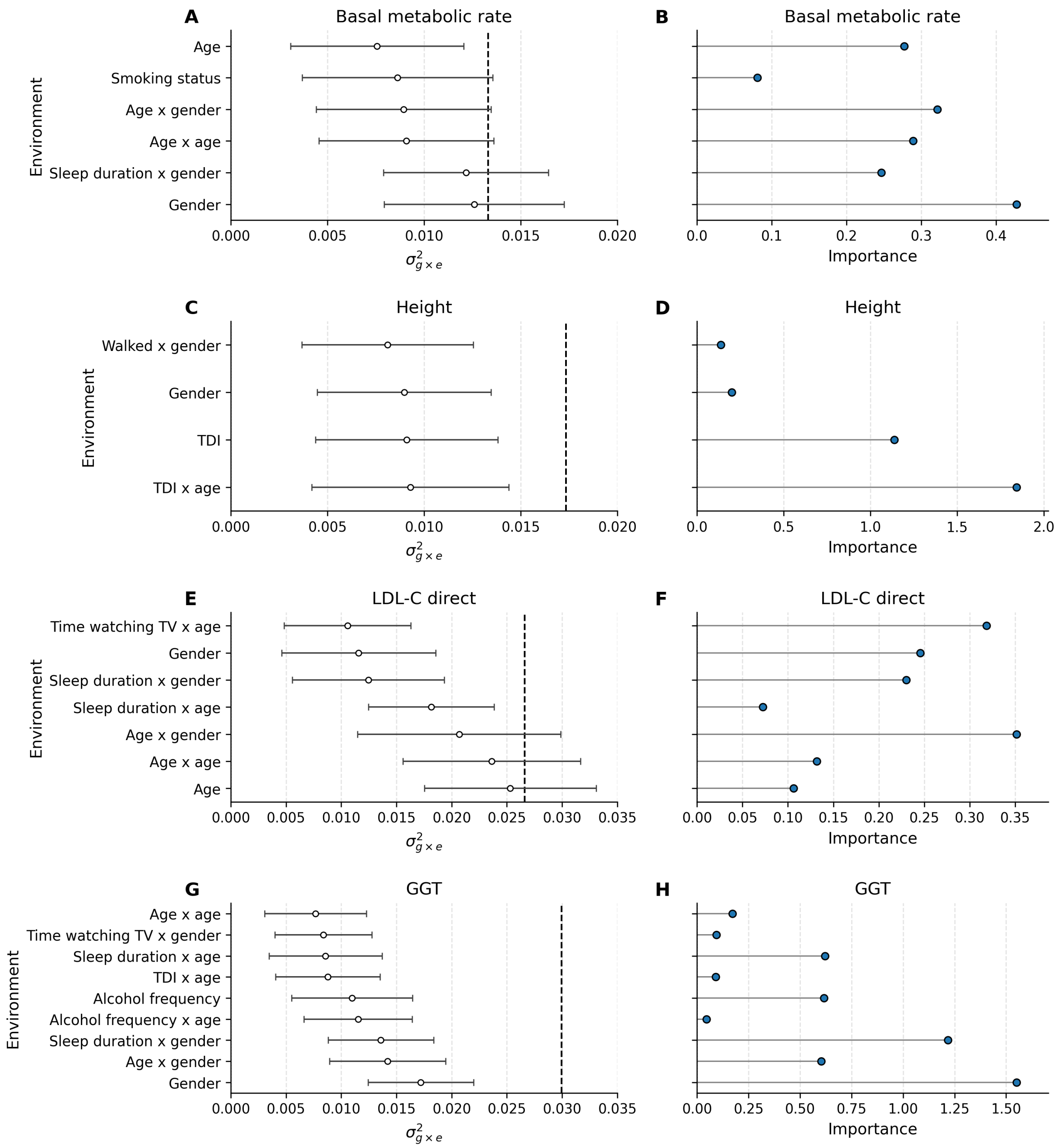


Figure S3: **Lifestyle G×E architecture across traits. A, C, E, G. Single-environment G×E estimates.** For each trait (basal metabolic rate, standing height, LDL-C direct, and GGT, respectively), we plot the estimated G×E variance components $\sigma^2_{g\times e}$ for each selected lifestyle exposure, with points denoting the estimates and horizontal bars indicating the standard errors. The dashed vertical line marks the G×E estimate for the learned multi-exposure embedding for that trait by ENGINE. **B, D, F, H. Learned embedding weights.** Absolute values of the learned embedding coefficients for the same set of exposures.

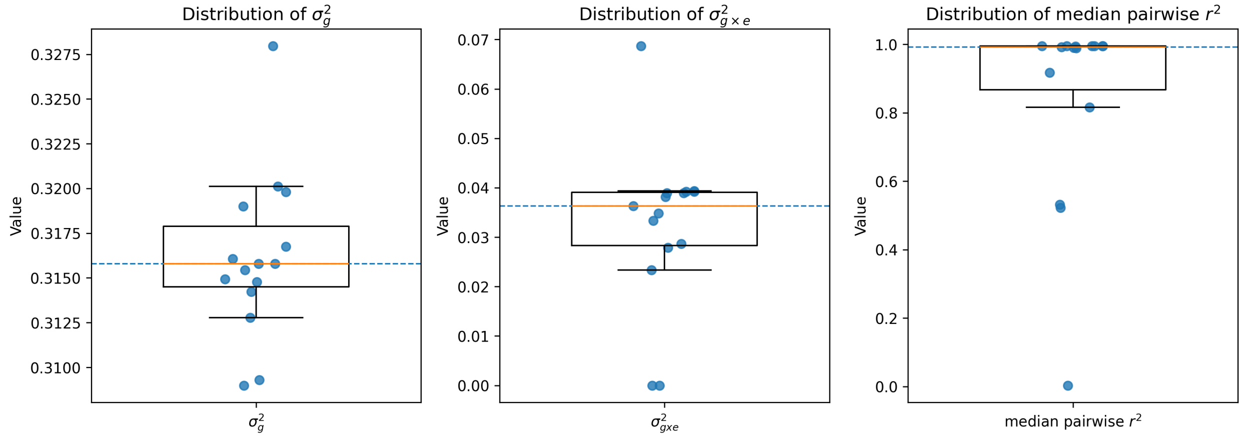


Figure S4: **Robustness of ENGINE across random seeds.** We repeated the BMI analysis with the ten selected environments, initializing ENGINE with 15 different random seeds. Across runs, we report the estimated $\sigma_g^2$, $\sigma_{g\times e}^2$, and the median pairwise $r^2$ between learned environmental embeddings.

### Supplemental Tables

Table S1: All notation.

| Symbol | Description |
|---|---|
| **Setup** | |
| $N$ | Number of individuals. |
| $M$ | Number of variants / SNPs. |
| $L$ | Number of measured environments (exposures). |
| $\boldsymbol{X} \in \mathbb{R}^{N\times M}$ | Standardized genotype matrix. |
| $\boldsymbol{y} \in \mathbb{R}^{N}$ | Standardized phenotype vector. |
| $\boldsymbol{E} \in \mathbb{R}^{N\times L}$ | Environment matrix; each column centered to mean 0 and scaled to unit variance. |
| $\boldsymbol{\alpha} \in \mathbb{R}^{L}$ | Learnable environment weight vector; constrained to $\\|\boldsymbol{\alpha}\\|_2 = 1$. |
| $\boldsymbol{e} = \boldsymbol{E}\boldsymbol{\alpha} \in \mathbb{R}^{N}$ | Environment embedding. |
| **Variance components** | |
| $\sigma_g^2$ | Additive (genetic) variance component. |
| $\sigma_{g\times e}^2$ | Gene-by-embedding interaction variance component. |
| $\sigma_{n\times e}^2$ | Noise-by-embedding (heteroskedastic) variance component. |
| $\sigma_n^2$ | Residual noise variance component. |
| $\boldsymbol{\sigma}^2$ | Variance components vector $(\sigma_g^2, \sigma_{g\times e}^2, \sigma_{n\times e}^2, \sigma_n^2)$. |
| **Random effects model (Eq. 1)** | |
| $\boldsymbol{\beta} \in \mathbb{R}^{M}$ | Additive genetic effects (random SNP effects). |
| $\boldsymbol{\gamma} \in \mathbb{R}^{M}$ | G×E interaction SNP effects along embedding $\boldsymbol{e}$. |
| $\boldsymbol{\delta} \in \mathbb{R}^{N}$ | Noise-by-environment (heteroskedastic) random effects; enters as $\mathsf{diag}(\boldsymbol{e})\,\boldsymbol{\delta}$. |
| $\boldsymbol{\epsilon} \in \mathbb{R}^{N}$ | Residual noise. |
| **Kernels / covariance** $(\mathbb{R}^{N\times N})$ | |
| $\boldsymbol{\Sigma}_y$ | Model covariance $\mathrm{Cov}[\boldsymbol{y}]$. |
| $\boldsymbol{K}_g = \frac{1}{M}\boldsymbol{X}\boldsymbol{X}^\top$ | Genetic relatedness matrix (GRM). |
| $\boldsymbol{K}_{g\times e}(\boldsymbol{\alpha}) = \frac{1}{M}(\boldsymbol{X}\odot\boldsymbol{e})(\boldsymbol{X}\odot\boldsymbol{e})^\top$ | G×E kernel (GRM after scaling rows by $\boldsymbol{e}$). |
| $\boldsymbol{K}_{n\times e}(\boldsymbol{\alpha}) = \mathsf{diag}(\boldsymbol{e})^2$ | N×E kernel (diagonal with entries $e_i^2$). |
| $\widetilde{\boldsymbol{\Sigma}}_y = \sum_\ell \widehat{\sigma}_\ell^2 \boldsymbol{K}_\ell$ | Plug-in covariance approximation used for SEs. |
| **MoM linear system (Eq. 2)** | |
| $\mathcal{L}(\boldsymbol{\alpha}, \boldsymbol{\sigma}^2)$ | Frobenius MoM objective $\left\\|\boldsymbol{y}\boldsymbol{y}^\top - \sum_r \sigma_r^2 \boldsymbol{K}_r\right\\|_F^2$. |
| $\boldsymbol{A} \in \mathbb{R}^{4\times 4}$ | Normal matrix with $A_{rs} = \mathrm{tr}(\boldsymbol{K}_r\boldsymbol{K}_s)$. |
| $\boldsymbol{b} \in \mathbb{R}^{4}$ | RHS with $b_r = \boldsymbol{y}^\top\boldsymbol{K}_r\boldsymbol{y}$. |
| $\widehat{\boldsymbol{\sigma}}^2$ | MoM estimate solving $\boldsymbol{A}\widehat{\boldsymbol{\sigma}}^2 = \boldsymbol{b}$. |
| **Optimization over $\boldsymbol{\alpha}$** | |
| $\boldsymbol{G}_y \in \mathbb{R}^{L\times L}$ | $\frac{1}{M}\big(\boldsymbol{X}^\top(\boldsymbol{E}\odot\boldsymbol{y})\big)^\top\big(\boldsymbol{X}^\top(\boldsymbol{E}\odot\boldsymbol{y})\big)$. |
| $\boldsymbol{G}_E \in \mathbb{R}^{L\times L}$ | $\frac{1}{M}\big(\boldsymbol{X}^\top\boldsymbol{E}\big)^\top\big(\boldsymbol{X}^\top\boldsymbol{E}\big)$. |
| $\boldsymbol{s}_y = \boldsymbol{y}\odot\boldsymbol{y} \in \mathbb{R}^{N}$ | Elementwise square of phenotype. |
| $\boldsymbol{s}_g = \mathsf{diag}(\boldsymbol{K}_g) \in \mathbb{R}^{N}$ | Diagonal of GRM. |
| $\boldsymbol{g} \in \mathbb{R}^{L}$ | Euclidean gradient of $\mathcal{L}$ w.r.t. $\boldsymbol{\alpha}$. |
| $\boldsymbol{g}_{23} \in \mathbb{R}^{L}$ | Trace-gradient contribution from terms involving $\mathrm{tr}(\boldsymbol{K}_{g\times e}^2)$ and $\mathrm{tr}(\boldsymbol{K}_g\boldsymbol{K}_{g\times e})$. |
| $\boldsymbol{g}_\top = \boldsymbol{g} - (\boldsymbol{\alpha}^\top\boldsymbol{g})\boldsymbol{\alpha}$ | Tangent-space projection on $\mathbb{S}^{L-1}$. |
| $\theta_{\max}$ | Maximum rotation angle (or step size). |

*Continued on next page*

**Table S1 (continued).**

| **Symbol** | **Description** |
|---|---|
| $\varepsilon_{\mathrm{ang}}, \varepsilon_{\mathrm{obj}}$ | Stopping tolerances (angular change; relative objective decrease). |
| **Sketching** | |
| $B$ | Number of probe vectors. |
| $\boldsymbol{W} \in \mathbb{R}^{B\times N}$ | Probe matrix; rows i.i.d. $\mathcal{N}(0, \boldsymbol{I}_N)$. |
| $\boldsymbol{U}_r = \boldsymbol{K}_r \boldsymbol{W}^\top$ | Kernel sketch for $\boldsymbol{K}_r$ (used for trace estimation). |
| $\boldsymbol{U}_g = \boldsymbol{K}_g \boldsymbol{W}^\top$ | Additive kernel sketch. |
| $\boldsymbol{Z}_\ell = \boldsymbol{X} \odot \boldsymbol{E}_{:,\ell}$ | Genotypes modulated by environment $\ell$. |
| $\boldsymbol{U}_{g\times e}[\ell, k]$ | Cached pairwise sketch. |
| $\boldsymbol{U}_{g\times e}(\boldsymbol{\alpha})$ | Assembled sketch of $\boldsymbol{K}_{g\times e}(\boldsymbol{\alpha})$ from $\{\boldsymbol{U}_{g\times e}[\ell, k]\}$. |
| $\boldsymbol{s}^{(1)}$, $\boldsymbol{s}^{(2)} \in \mathbb{R}^{L\times L}$ | Contractions used to assemble $\boldsymbol{g}_{23}$. |
| $\boldsymbol{C} \in \mathbb{R}^{L\times L}$ | Upper-triangular pair-weight matrix used conceptually for $\boldsymbol{g}_{23} = (\boldsymbol{C} + \boldsymbol{C}^\top)\boldsymbol{\alpha}$. |
| **Regularized cross-fitting** | |
| $\boldsymbol{X}_{(1)}, \boldsymbol{X}_{(2)} \in \mathbb{R}^{N\times(M/2)}$ | Variant split for cross-fitting (disjoint SNP halves). |
| $\boldsymbol{\alpha}_{(1)}, \boldsymbol{\alpha}_{(2)}$ | Embeddings learned on split halves. |
| $\widehat{\boldsymbol{\sigma}}^2_{(1)}, \widehat{\boldsymbol{\sigma}}^2_{(2)}$ | Variance components estimated on split halves. |
| $\lambda$ | $\ell_1$ regularization strength for $\\|\boldsymbol{\alpha}\\|_1$. |
| $\tau$ | Step length used in soft-thresholding update (followed by renormalization). |

Table S2: UK Biobank field IDs for environmental exposures and phenotypes used in this study.

| Category | Variable | UKB field ID |
|---|---|---|
| Environment / non-dietary | Sex | 31 |
| Environment / non-dietary | Townsend deprivation index at recruitment | 189 |
| Environment / non-dietary | Days/week walked 10+ minutes | 864 |
| Environment / non-dietary | Days/week of moderate physical activity (10+ minutes) | 884 |
| Environment / non-dietary | Days/week of vigorous physical activity (10+ minutes) | 904 |
| Environment / non-dietary | Time spent watching television (TV) | 1070 |
| Environment / non-dietary | Sleep duration | 1160 |
| Environment / dietary | Cooked vegetable intake | 1289 |
| Environment / dietary | Oily fish intake | 1329 |
| Environment / dietary | Non-oily fish intake | 1339 |
| Environment / dietary | Processed meat intake | 1349 |
| Environment / dietary | Poultry intake | 1359 |
| Environment / dietary | Beef intake | 1369 |
| Environment / dietary | Lamb/mutton intake | 1379 |
| Environment / dietary | Pork intake | 1389 |
| Environment / dietary | Cheese intake | 1408 |
| Environment / dietary | Salt added to food | 1478 |
| Environment / dietary | Tea intake | 1488 |
| Environment / non-dietary | Alcohol intake frequency | 1558 |
| Environment / non-dietary | Age at recruitment | 21003 |
| Environment / non-dietary | Smoking status | 20116 |
| Phenotype | Standing height | 50 |
| Phenotype | Body mass index (BMI) | 21001 |
| Phenotype | Basal metabolic rate | 23105 |
| Phenotype | Gamma glutamyltransferase (GGT) | 30730 |
| Phenotype | LDL cholesterol (direct) | 30780 |

Table S3: **ENGINE optimization and sketching hyperparameters for the biobank-scale run.** Angular hyperparameters are reported in radians: the recommended settings correspond to $\theta_{\max} = 25° \approx 0.44$ and $\varepsilon_{\text{ang}} = 0.05° \approx 8.73 \times 10^{-4}$.

| **Parameter** | **Value** |
|---|---|
| $N$ (individuals) | 291,273 |
| $M$ (SNPs) | 454,207 |
| $L$ (environments) | 8 |
| Seed | 42 |
| $B$ (Hutchinson probes) | 100 |
| Maximum iterations | 2000 |
| Step size ($\eta$) | 1 |
| Maximum rotation ($\theta_{\max}$; radians) | 0.44 |
| Angular tolerance ($\varepsilon_{\text{ang}}$; radians) | $8.73 \times 10^{-4}$ |
| Objective tolerance ($\varepsilon_{\text{obj}}$) | $10^{-6}$ |
